# Vasculature segmentation in 3D hierarchical phase-contrast tomography images of human kidneys

**DOI:** 10.1101/2024.08.25.609595

**Authors:** Yashvardhan Jain, Claire L. Walsh, Ekin Yagis, Shahab Aslani, Sonal Nandanwar, Yang Zhou, Juhyung Ha, Katherine S. Gustilo, Joseph Brunet, Shahrokh Rahmani, Paul Tafforeau, Alexandre Bellier, Griffin M. Weber, Peter D. Lee, Katy Börner

## Abstract

Efficient algorithms are needed to segment vasculature in new three-dimensional (3D) medical imaging datasets at scale for a wide range of research and clinical applications. Manual segmentation of vessels in images is time-consuming and expensive. Computational approaches are more scalable but have limitations in accuracy. We organized a global machine learning competition, engaging 1,401 participants, to help develop new deep learning methods for 3D blood vessel segmentation. This paper presents a detailed analysis of the top-performing solutions using manually curated 3D Hierarchical Phase-Contrast Tomography datasets of the human kidney, focusing on the segmentation accuracy and morphological analysis, thereby establishing a benchmark for future studies in blood vessel segmentation within phase-contrast tomography imaging.

## Introduction

Because blood vasculature extends to all organs of the body, identifying vessels in medical images (“vessel segmentation”) is an important part of image processing in applications ranging from basic science research to clinical care.

As the vascular network is an inherently 3D and multi-scale spanning structure, imaging techniques that do not require physical subsampling to achieve high-resolution images across multiple scales are the ideal tool for vascular analysis^1^; Hierarchical Phase-Contrast Tomography (HiP-CT) is such a technique^2^. Leveraging the European Synchrotron Radiation Facility’s (ESRF) Extremely Brilliant Source (EBS), the world’s first high-energy fourth generation synchrotron source, HiP-CT enables imaging of intact human organs at unprecedented scale and resolution. HiP-CT can be used to map and quantify the arterial vascular network of an intact human kidney down to the arteriolar level^3^. However, the creation of 3D models from HiP-CT data using traditional manual segmentation techniques is very time-consuming and labor-intensive. This labor-cost barrier impedes the broader utilization of 3D data to answer questions proposed by the biomedical community, including understanding exosome transport.

Considering such aims and challenges, a global citizen science competition was organized on Google’s machine learning (ML) platform, Kaggle, to develop open-source machine learning solutions capable of automatically segmenting large datasets to generate 3D models of the blood vasculature of the human kidney. The Kaggle platform has been previously leveraged to collaboratively develop machine learning solutions for complex biomedical tasks thereby advancing the field^4–7^. The competition presented in this paper focuses on 3D data, challenging teams to develop solutions to accurately segment blood vessels in HiP-CT images to create high quality 3D models of the vessels from organs such as the kidney. Such segmented 3D models can be used to obtain additional information such as length, diameter, branching angles, tortuosity, inter-vessel distance, etc.

The two sponsors of this competition are the NIH Common Fund’s Human BioMolecular Atlas Program (HuBMAP) and Cellular Senescence Network (SenNet) Program. Both provide examples of how researchers can benefit from improved methods of vessel segmentation and served as motivation for the competition:

1. HuBMAP is constructing a map of all the cells in the healthy adult human body. It envisions blood vasculature pathways serving as landmarks to describe the location of cells in organs and tissue^8–10^. In this “Vasculature Common Coordinate Framework” (VCCF)^11^, accurate vessel segmentation is essential to define these landmarks. As part of this work, HuBMAP has published an initial database of more than 900 blood vessels based on literature review and domain expert curation^12^. However, this needs to be linked to experimental data for validation and to extend it to microvasculature, where there are many gaps in existing knowledge.
2. SenNet studies senescent cells of the human body—in health and in disease—across the lifespan^13,14^. A key challenge is understanding the secretome, i.e., the set of proteins these cells secrete into the extracellular space and/or the bloodstream. Exosomes—a subcategory of secretomes—are membrane-bound extracellular vesicles used to transport diverse cargo to neighboring cells and to other organs. However, understanding the generation, transportation, and ingestion of exosomes requires data on the human blood vasculature—the main exosome transport system. By analyzing this complex highway of blood flow which includes determining the pathway, throughput/vessel size, and 3D orientation in space of the blood vasculature, we aim to contribute to the understanding of exosome diffusion via the bloodstream, how exosomes can play roles in organ-to-organ communication, and where exosomes might aggregate due to branching or tortuosity of the vessels—analogous to river sediments.^15–17^

This paper presents the competition setup, the curated HiP-CT 3D datasets, the competition metric, and the top-5 winning solutions. Additionally, to enable a holistic evaluation of ML model performance, three additional analyses are provided: quantitative (based on additional metrics), qualitative (based on visual analysis of predictions), and morphological (based on vasculature morphology features). Due to the novelty of HiP-CT data and lack of gold standard segmentations for vasculature in such data, the paper presents a benchmark dataset, performance metrics, and benchmark scores for future research in the domain, which can accelerate the augmentation of our knowledge of the blood vasculature system in support of answering key questions posed by the research community.

## Results

### Competition design

The “*SenNet + HOA: Hacking the Human Vasculature in 3D”* competition—running from November 7, 2023 through January 30, 2024—aimed to develop machine learning solutions for segmentation of blood vasculature in 3D HiP-CT scans of the human kidney. The imaging datasets were collected as part of the Human Organ Atlas effort (https://human-organ-atlas.esrf.eu). Vessel segmentation is a challenging task for various reasons: it is an imbalanced class problem (where prediction targets are disproportionately lower in volume compared to non-target background), there is a very low tolerance for error since small losses in segmentation connectivity can lead to large variations in simulated function^18^, there can be collapse or infilling of vessels during preparation which makes them challenging to segment^19^, there is wide image variability due to natural human anatomical variation and due to the nature of imaging post-mortem organs, and there exist relatively small training datasets due to the novelty of the HiP-CT imaging technique. These issues were documented in previous work by the authors^19^ which also showcased the performance of a baseline NN-Unet^20^ model on HiP-CT data. A key challenge for participants was to develop solutions that can handle large 3D datasets and that generalize to different datasets containing variability in resolutions, image intensities, and donors.

The HiP-CT datasets used in the competition were sourced from the kidneys of five adult donors. **Table 1** provides an overview of the donor demographics, image and scanning metadata and gold standard label metadata associated with each dataset (Full scanning and donor medical metadata can be found at https://human-organ-atlas.esrf.eu, DOI and link for each dataset are provided in **Supplementary Table 1**). **Figure 1** gives an overview of the competition data and setup. The five 3D image volumes were split into training and test datasets. The training dataset contained four image volumes, including three whole kidney (∼50µm/voxel) datasets and one high resolution hierarchical Volume-of-Interest (VOI) dataset. Two of these whole kidney datasets contained a mix of sparse labels as well as dense labels (see **Methods**). The test set was further divided into a public test set and a private test set. The public test set contained one partial kidney volume with 50.28 µm/voxel data (1,013 slices). The private test set contained one partial kidney volume with 63.08 µm/voxel data (501 image slices), see **Methods** for details. It should be noted that the public test dataset was very similar to the training dataset e.g. the same donor as kidney 1 (but left rather than right kidney) and collected on the same beamline BM05. By comparison the private test data was from a different donor and collected with the same technique but on a different beamline, BM18. This choice of test data was made to reflect the real challenges faced by our research teams e.g., to create models from limited training datasets such that the models generalize effectively to new donors and are robust to some changes in the imaging setup. In total, approximately 600 human hours were spent in segmentation and verification of the gold standard labels for the competition dataset, underscoring the time and resource consuming nature of the segmentation problem and the need for automated methods for the task.

**Table 1.** Competition dataset details, including metadata and donor demographics. While a Dataset 4 (without gold standard labels) was originally planned for inclusion in the competition dataset, it was removed before competition launch.

| Kaggle Donor ID | Donor ID | Segmentation labels | Voxel size (μm) | Image z-slices | Dataset usage | Beamline | Sex | Age |
| --- | --- | --- | --- | --- | --- | --- | --- | --- |
| 1 | LADAF 2021-17-Right-Kidney | Whole kidney densely labeled | 50 | 2,279 | Training | BM05 | M | 63 |
| 1 | LADAF 2021-17-Right-Kidney | Whole VOI densely labeled | 5.2 | 1,397 | Training | BM05 | M | 63 |
| 2 | S-20-28-Kidney | Whole kidney sparsely labeled (65% of vessels) | 50 | 2,217 | Training | BM05 | M | 84 |
| 3 | LADAF-2020-27-Kidney-2 | Whole kidney part densely, remainder sparsely labeled, (85% of vessels) | 50.16 | 501 dense<br>1,035 sparse | Training | BM05 | F | 94 |
| 5 | LADAF-2021-17-Left-Kidney | Partial whole kidney densely labeled | 50.28 | 1,012 | Public test | BM05 | M | 63 |
| 6 | LADAF-2022-13-Bottom-Kidney | Partial whole kidney densely labeled | 63.08 | 501 | Private test | BM18 | M | 85 |

**Figure 1.**
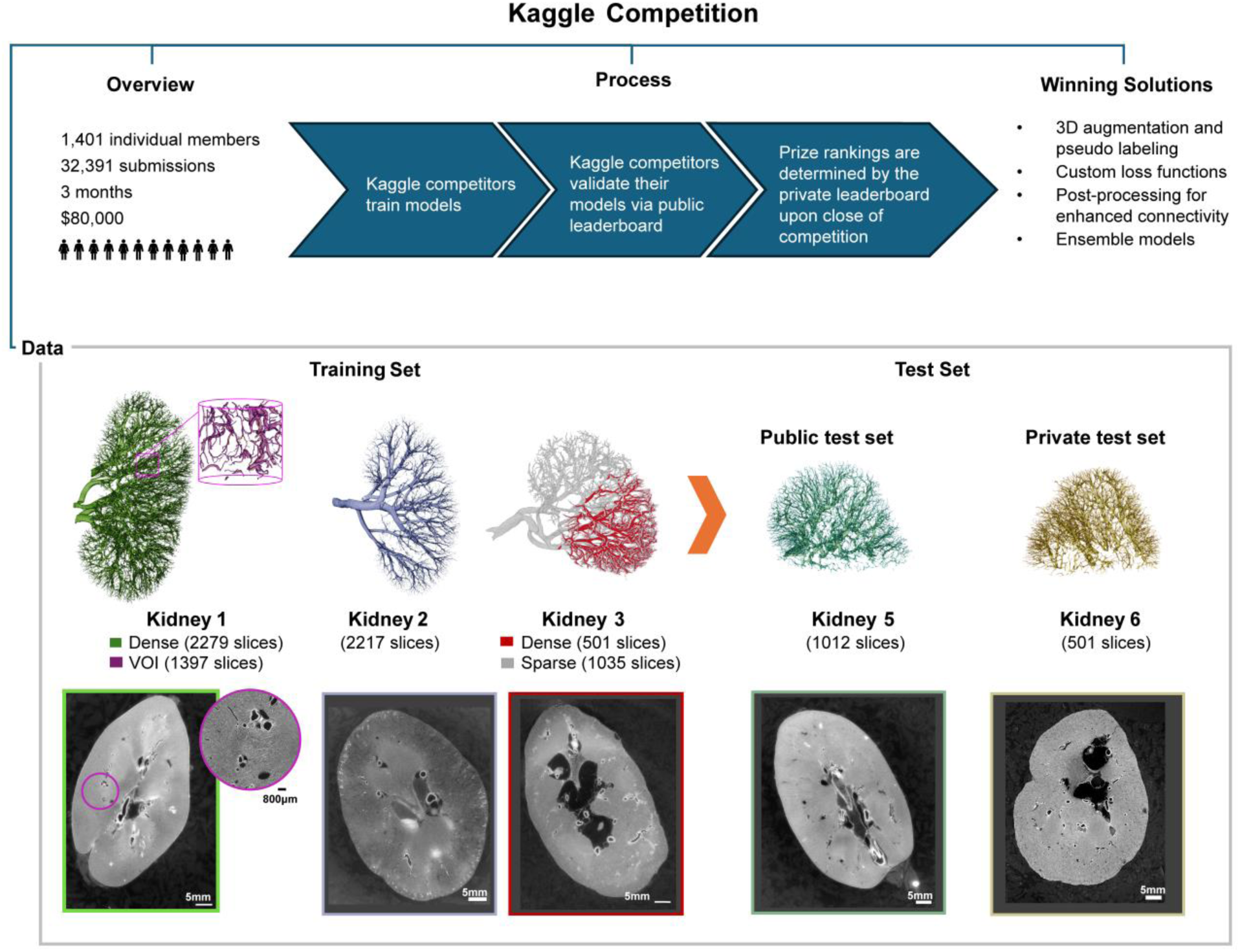
An overview of the competition setup and process, including 3D renderings of the gold standard labels and representative 2D slices for all data in the competition.

The challenges associated with the creation of high quality gold standard labels for training ML models is well known^21^. However, with new modalities such as HiP-CT, this poses a particular challenge as there is no large pool of potential expert curators who are familiar with the data type and the particular artifacts that can be present. In order to acknowledge the challenge with creating this gold standard, we chose to create a labeling protocol involving three independent curators (see **Methods** for details). Using this protocol, we were able to provide both dense labeling (i.e., where the third curator found 100% of vessels were labeled), and to provide sparse labeling with an estimate of the sparsity. This allowed some teams, e.g., Team 3, to use the sparse labels effectively as training data within a pseudo-labeling context. This was critical in the competition design as it is relatively fast for an experienced curator to sparsely label an image volume but highly time consuming to label it densely.

In addition, a high-resolution VOI was provided, which is a subset of the lower resolution whole kidney training dataset that can be rigidly registered to the whole kidney dataset to provide more accurate labels for smaller vessels. Previously, we have shown how these higher resolution VOIs can be used to validate manual segmentation in the lower resolution dataset^3^. Interestingly, none of the teams attempted to utilize or extend this multi-resolution approach to improve the segmentation of small vessels, although teams that used this dataset as additional training data, such as Team 1, found a greater proportion of the smaller vessels.

In addition to the competition data, participants were allowed to use publicly available external datasets. Some teams used other HiP-CT datasets (gold standard labels absent) available via the Human Organ Atlas data portal (https://human-organ-atlas.esrf.eu) to help improve the performance of their solutions. All inference code was submitted via the Kaggle submission portal and was run on the public and private test sets within the Kaggle infrastructure, leading to team rankings on the public and private leaderboards (LB), respectively (see **Methods**). Throughout the competition, the private LB scores and rankings are not visible to the participants which forces them to use the public LB scores and rankings as a validation dataset for their experiments. The test datasets are not visible to the participants but they are provided 3 z-slices each from both test sets as examples. They are also provided the voxel resolution for both test sets.

Algorithm performance was judged based on the Normalized Surface Dice (NSD) metric as previously proposed by the Google Deepmind team^22,23^, with tolerance threshold set to 0 (see **Methods**). At the end of the competition, the top-5 teams on the final private leaderboard won the performance prizes and the associated prize money (see **Methods**).

The competition observed participation by 1,401 individual members divided across 1,149 teams from 78 countries. There were 204 members that participated for the first time in a Kaggle competition, five of which were in the top-100 on the final private leaderboard. There were 32,391 unique code submissions, 500 public notebooks created, 756 discussion comments, and 200 discussion forum topics. There was also a separate Discord server created for the competition where teams engaged in more informal discussions throughout the competition, although this was not monitored by the hosts/authors. **Figure 2a** graphs the changes in the leaderboard scores, forum activity, and number of participants during the three-month period of the competition. **Figure 2b** shows the number of submissions versus the highest private score for all teams.

**Figure 2.**
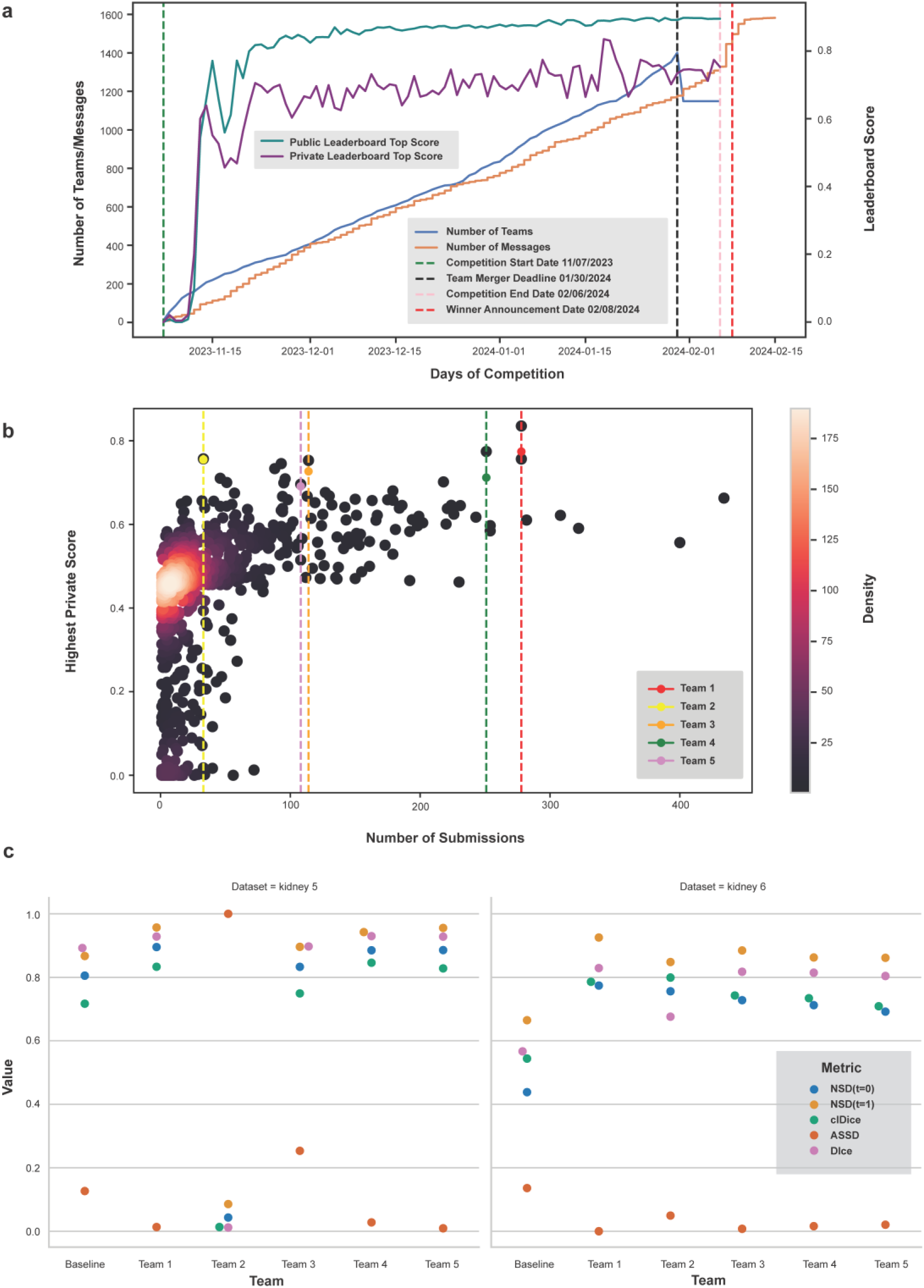
**a.** Number of teams and messages, and leaderboard high scores per day over competition period. **b.** Number of submissions vs. highest private leaderboard score for each of the 1,149 teams as a heatscatter. **c.** Scores for competition metric and additional metrics for top-5 teams on both test sets. ASSD is normalized between 0-1 for visualization (lower is better).

### Overview of winning solutions

In this competition, in general, teams employed a variety of techniques to tackle the challenge, including customized U-Net architectures, different data augmentation techniques, and complex loss functions. Notably, methods integrating data augmentation and self-training paradigms emerged as effective solutions to address challenges related to the small dataset. Teams utilized pseudo-labelling techniques, either refining pseudo labels from sparse to dense iteratively, or creating labels for external datasets. Additionally, post-processing steps such as to eliminate disconnected vessels also helped improve solutions.

The competition highlighted that the U-Net architecture^24^ remains a strong base model for medical image segmentation. All of the top five teams used variants of U-Net—2D, 2.5D, or 3D—with different backbones, customizing the models to enhance performance while keeping the U-shaped structure unchanged. Among the advanced backbones, MaxViT-Large-512^25^ stood out for its ability to handle large input sizes (like 512×512 pixels), making it suitable for high-resolution HiP-CT images. By leveraging the dual capabilities of Convolutional Neural Networks^26^ (CNN) and Transformers^27^, MaxViT-Large-512 enhanced U-Net’s ability to accurately segment complex anatomical structures. Additionally, ensembling different U-Net architectures was popular in the competition, with three out of the top-5 winning teams utilizing ensemble models. Individual models might overfit to specific patterns in the training data; ensembling reduces the risk of overfitting by averaging out biases from individual models, resulting in better generalization on unseen data. Additionally, by combining the strengths of different models, an ensemble provides more robust predictions, effectively handling anomalies and noise in the data. Larger sliding-window size generally had better performance as it was able to remove more artifacts at the window boundary introduced by sliding-window inference.

Team 1 developed a tailored 2.5D U-Net^24^ with ConvNeXt-tiny^28^ as the encoder, using a combination of loss functions to directly optimize for NSD. Team 2 utilized a 3D U-Net^29^, focusing on improving generalization through extensive data augmentation and morphological post-processing. Team 3 combined multiple U-Net-based architectures with CNN and Transformer models, refining sparse labels with dense ones for better segmentation performance. Team 4 combined 2D and 3D models, leveraging test-time augmentation (TTA) and pseudo-labeling to enhance accuracy. Finally, Team 5 ensembled three 2D U-Net models, integrating pseudo-labeling and boundary loss to sharpen boundary predictions. Through innovative data preprocessing, careful model selection and hyperparameter choice, and effective post-processing, these teams achieved high accuracy, securing top-five positions in the competition.

Furthermore, there were some key implementation details in the solutions proposed by the top-5 teams that highlighted differences in their approaches. Team 1: Defining a custom Marching Cube Loss function, adding an extra stem block to the model to leverage high resolution information, using 3D rotations, and generating more data by slicing along different axes. Team 2: Generating more data using 3D affine transformations on training sets, and removing distantly placed disconnected chunks to reduce false positives. Team 3: Refining pseudo labels from sparse to dense iteratively, emulating test set magnification, and intensity augmentation. Team 4: Percentile normalization of training data, pseudo labels on external data, Boundary Difference over Union loss for both 2D and 3D models, and sliding window inference. Team 5: Creating soft-labels with pseudo labels on external data, combined with composite loss with more weight to the boundaries of masks, and 3D interpolation to account for the different resolutions.

As discussed earlier, vessel segmentation is generally a difficult image processing challenge, due to the multi-scale nature and small structure of vessels. With HiP-CT data of human organs, several additional challenges are present, all of which are compounded by a low tolerance for connectivity error in the final segmentations^19^. Most of the teams attempted to overcome the issue of a small training dataset by employing different augmentation techniques and pseudo-labeling approaches. For example, Teams 1 and 2 significantly enhanced their scores with the use of 3D rotational augmentation. After implementing this technique, Team 1’s private score increased from 0.682 to 0.835. Team-3’s strategy of iterative pseudo-labeling enabled training of models on an expanded dataset, securing it the third rank in the competition.

To tackle the issue of unbalanced classes, some teams introduced various boundary losses, which take the form of a distance metric based on contours rather than regions. This approach helped mitigate the challenges associated with highly unbalanced datasets by focusing on integrals over the interfaces between regions, rather than unbalanced integrals over the regions themselves. For instance, Team 1 experienced a notable improvement after integrating a combination of focal, dice, and boundary losses with a custom loss function (see **Supplementary Note 1**), elevating its private score from 0.534 to 0.682. Meanwhile, Team 4’s use of BoundaryDOULoss^30^ (see **Supplementary Note 4**) boosted its public LB score by 1.5% and its private LB score by 5%. To address the connectivity issue, Team 2 effectively used a post-processing step to enhance connectivity and eliminate false positives, by removing small unconnected segments with depth-first search. In order to mitigate the effect of variability of the imaging data, Team 3 emulated a test set magnification to have matching resolutions. Detailed technical information for each winning team’s solution, including information for some other teams, is provided in **Supplementary Notes 1-6**.

### Quantitative analysis of solutions

The top-winning team, Team 1, reached a NSD score of 0.774 on the final private LB and also ranked first on the public LB with a score of 0.898, followed by 0.755 for Team 2, 0.727 for Team 3, 0.712 for Team 4, and 0.691 for Team 5. The top-5 teams made a total of 784 out of 32,391 submissions (2.42% of total submissions). **Supplementary Table 2** lists public and private LB scores for all five winning teams, including the baseline NN-Unet model’s performance, and a brief summary of their solutions.

In comparing team rankings across public and private LBs, Team 1’s solution proved to be the most stable as it ranked at the top on both LBs. Team 4 and Team 5 were fairly stable: Team 4 rising by one place and Team 5 rising by five places on the private LB compared to the public LB. Solution by Team 2 was the most unstable, jumping 1,022 places on the private LB, while Team 3 jumped by 568 places. Team 2’s solution was highly overfitted to the private test set, and scored 0.04 NSD score on the public LB.

Additionally, authors previously identified certain key metrics for the vessel segmentation problem after reviewing the existing literature^19^. These additional metrics—Dice Similarity Coefficient (Dice), Centerline Dice (clDice), NSD (tolerance=1), and Average Symmetric Surface Distance (ASSD)— are computed for the predictions of the top-5 winning teams (see **Methods** for details on the metrics). All computed metrics for both test sets are provided in the **Supplementary Table 3** and visualized in **Figure 2c**. The difference between the NSD scores at tolerance 0 and tolerance 1 are larger for the private test set than for the public test set across all teams. For the private test set, Team 1 scores higher than the other 4 teams across all additional metrics, except in the case of clDice where Team 2 performs slightly better. While the performance of Team 3 is comparable to other teams on the private test set, it performs worse on the public test set across multiple metrics, thereby highlighting the low generalizability of solutions by Team 2 and Team 3.

A fundamental challenge in organizing such competitions that need to be scored based on a single metric is that changing the metric can change the outcome of leaderboards as well as the ability of the created solutions^31,32^. In light of this, it is valuable to see the correlation between public and private LB rankings as well as the effect of rankings based on different metrics. Subsequently, Kendall’s Tau^33^ (KT, see statistical analysis in **Methods**) is computed and a value of 0.0253 (p-value=0.795) is found for rankings of private and public LB (top-50 teams). This highlights a high shake-up in rankings on both LBs, showing that, in general, solutions by top-50 participants had low generalizability. To check rank stability based on different metrics, KT is computed for all additional metrics with respect to the original competition metric. The KT for top-50 teams on the private test set based on NSD (tolerance=0) and clDice is 0.502 (p-value=2.6839e-07), KT for NSD at tolerance 0 vs. 1 is 0.472 (p-value=1.2773e-06), KT for NSD (tolerance=0) and Dice is 0.387 (p-value=7.0881e-05), and KT for NSD (tolerance=0) and ASSD is 0.3877 (p-value=7.0881e-05).

**Supplementary table 3** shows all metric scores for top-50 teams for both test sets.

### Qualitative analysis of winning solutions

Due to the low generalizability of solutions by Team 2 and Team 3, the qualitative and morphological analysis focuses on Teams 1,4 and 5. **Figure 3** shows a Maximum Intensity Projection (MIP) for the gold standard labels for the private test set and the predicted labels from Teams 1,4, and 5. Similar figures for Team 2 and Team 3 are presented in **Supplementary Figure 1**. Visual analysis across the three teams (Team 1, 4, and 5), shows a high degree of missing vessel connectivity, particularly for the small vessels which can be seen in the insets in **Figure 3** (green arrows). This unconnected vessel morphology is particularly evident at the outer edge of the kidney where the majority of vessels are thinner and the variations between the teams are more visually pronounced. Interestingly, Team 4’s solution appears to have the least number of small unconnected components, favoring larger vessel detection, whereas Team 5’s solution detects parts of the small vessels but they are highly disconnected. Team 5 also finds a large number of small unconnected components outside the boundary of the kidney itself (red inset in **Figure 3**). This is a feature which Team 4 were able to remove in post processing though a simple but effective masking of the kidney outer edge, a strategy which dramatically reduces the number of false positive pixels. Team 1’s solution visually shows better connection of the small vessels in comparison to Team 5’s potentially due to the extra stem in their model designed to capture these smaller vessels. Additionally, Team 1’s solution appears to detect tubular structures that are not in the gold standard (orange arrowhead in **Figure 3**). Several of these structures, when compared back to the raw data, appear to be correctly identified vessels that were not found by the manual segmentation process. This highlights that manual segmentation of data, even when performed by multiple curators, can only ever be considered as a gold standard (with some degree of such acceptable data noise) rather than a complete ground truth. Finally, all teams appear to predict vessels that are thinner than the gold standard labels.

**Figure 3.**
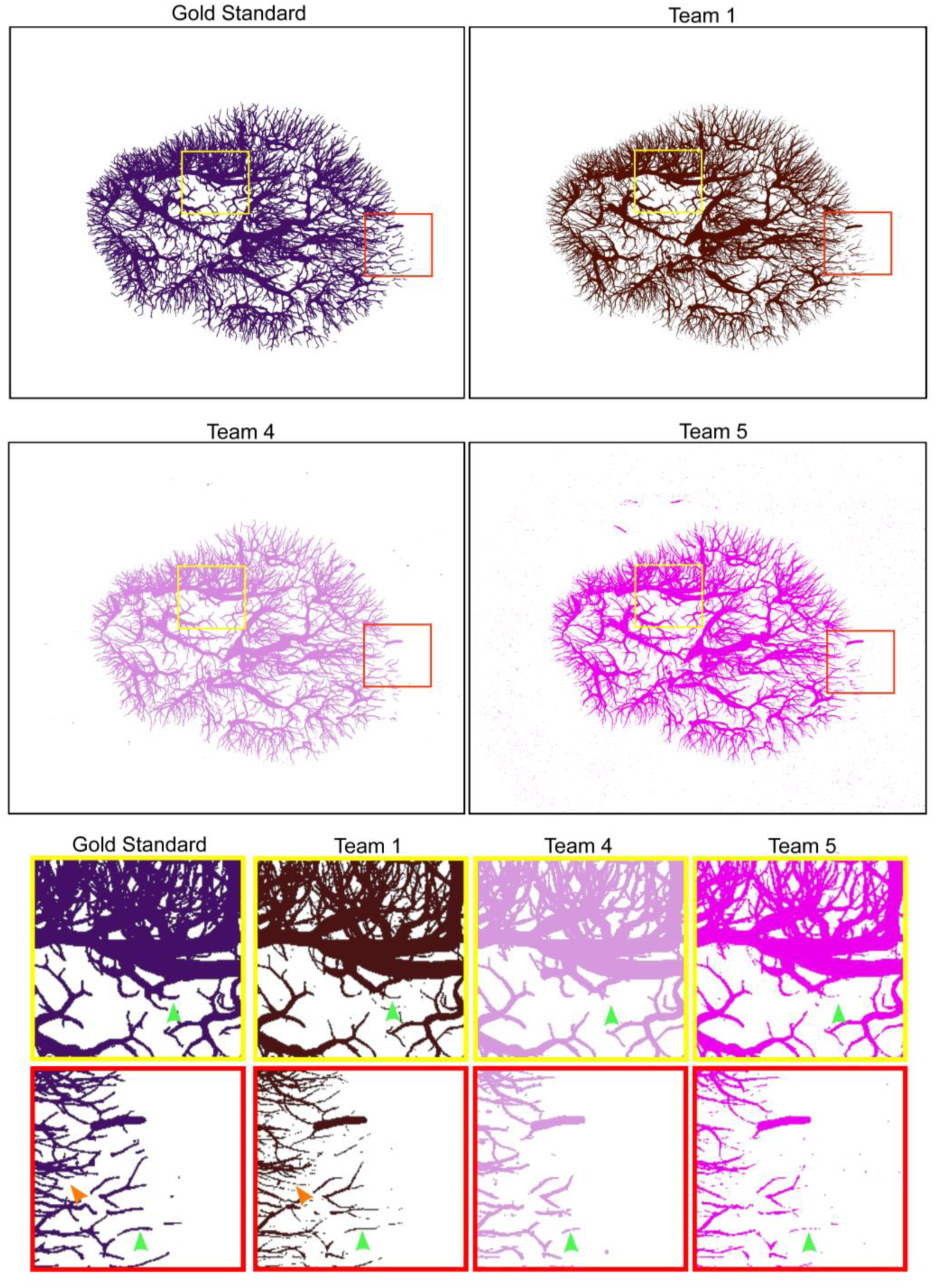
Maximum Intensity Projections (MIP) for the labels (private test set) for gold standard, Team 1 predictions, Team 4 predictions, and Team 5 predictions, with two insets per team in the yellow and red squares, respectively. Green arrows show the same location in each prediction highlighting the missing and unconnected vessels in Teams 1,4 and 5. The orange arrow in the gold standard and Team 1 show an instance of a vessel predicted by Team 1’s model that is not in the gold standard but is, in fact, in the raw data.

As the test datasets—both public and private—were only portions of the whole intact kidneys, we computed inference for each team’s model on the remaining parts of the private test set (kidney 6) (**Figure 4a**) and the public test set (kidney 5) (**Supplementary Figure 4).**The outputs in **Figure 4a** serve to further demonstrate the differences, particularly in the ability to capture larger vessels between these models and their generalizability. Team 1’s solution, which appeared to find many of the smaller vessels in test data, evidently misses large portions of the larger vessels in the whole private test data (kidney 6), and to a lesser extent in the public test data (Kidney 5). The larger vessels are evidently more fully captured by Teams 3-5 in both the public and private test datasets. The lack of generalizability in Team 2 is highly apparent when comparing the outputs for kidney 5 (**Supplementary Figure 4)** and kidney 6 (**Figure 4a)**.

**Figure 4.**
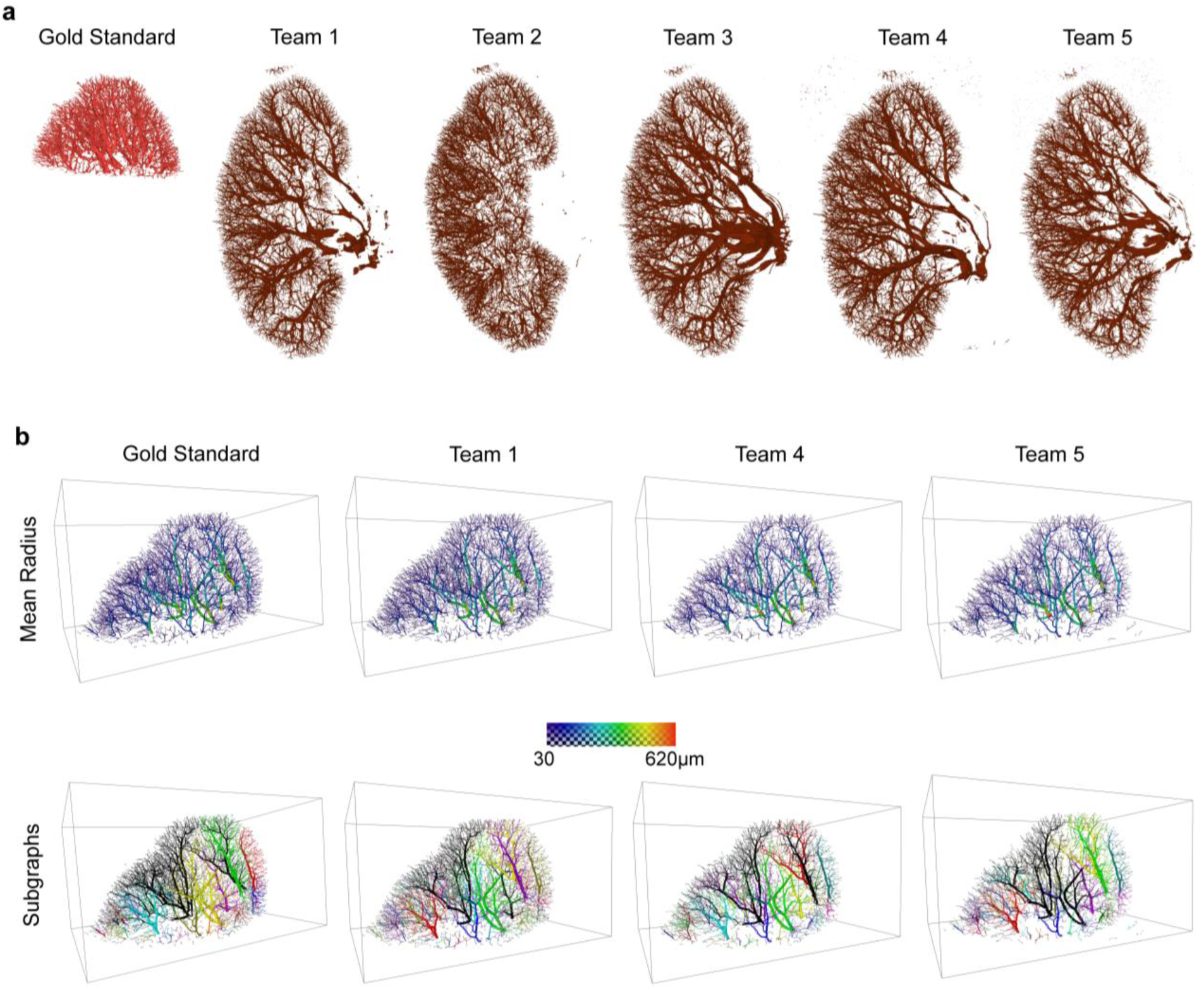
**a.** Visualization of the 3D output for inference of each team on the whole intact kidney datasets for the private test data (Kidney 6). Gold standard (red) shows the part of the whole kidney that was fully labeled and was part of the original competition dataset. **b.** Visualization of the skeleton forms of the vessel network for the gold standard and teams 1, 4 and 5 for the Private test data (Kidney 6). In each case, each vessel segment is rendered with the mean radius. In the first row the vessels are colored according to each segment’s mean radius. In the second row each unconnected subgraph is in a different color.

### Morphological analysis of segmentations

While the quantitative segmentation metrics and qualitative inspection of the voxelized output provides an insight into the performance of different solutions, a comparative morphological approach that aligns with the eventual intended use of the segmentations is highly relevant to assessing model biases and outputs. Given that our ultimate aim is to use the vascular segmentations to 1) model exosome transport, 2) extract vessel morphology metrics to create a VCCF; the segmented vascular networks will be skeletonized. Skeletonization reduces the 3D voxel representation to a skeleton representation described by segments and nodes—with each segment defined by start and end nodes—and having a length, radius, and connectivity to other segments (see **Methods** for further details). Here, we skeletonize each of the predicted segmentation outputs to investigate how the differences in segmentation from the various models lead to variation in the extracted vascular network geometry.

**Figure 4b** shows the spatial variations in the mean radius and subgraphs (unconnected portions of the network) between model outputs. While the number of subgraphs varies between teams, Teams 1,4, and 5 find the same larger vessel trees. Variations are mostly restricted to the smaller disconnected vessels at the kidney periphery which was also noted in the qualitative analysis. Similarly, while the spatial distribution of radii is similar, the absolute values vary between each solution and the gold standard. **Supplementary Figure 2** shows the same visualizations for Team 2 and Team 3. **Supplementary Figure 3** shows the skeleton forms of the vessel networks for all teams on public test data.

**Figure 5** highlights these morphological differences between the skeletonized networks geometry for the private test data (similar plots for public test data are provided in **Supplementary Figure 4**). For all teams, the number of subgraphs is higher than the gold standard, which indicates many short unconnected segments that form the partial path of the vessels (**Figure 5a**). The shorter mean length of segments **(Figure 5c**) also supports this conclusion as does the proportions of terminal nodes compared to branched nodes which is higher than the gold standard for all teams other than Team 2. The mean radius (**Figure 5b**) has large variability highlighting the challenge of accurately determining vessel thickness. In all cases, the mean radius is lower than in the gold standard which appeared to be the case in the qualitative analysis. The thinner vessels are also reflected by the much higher NSD score where there is a 1-voxel tolerance (see **Figure 2**). For tortuosity and branching angle, values closer to the gold standard and baseline models indicate better modeling of natural vessel shapes. As tortuosity is associated with length, it is interesting to note that while Team 4 has a lower vessel length than Teams 3 and 5, it has a higher tortuosity, indicating that many short but highly curved vessels are present in this case.

**Figure 5.**
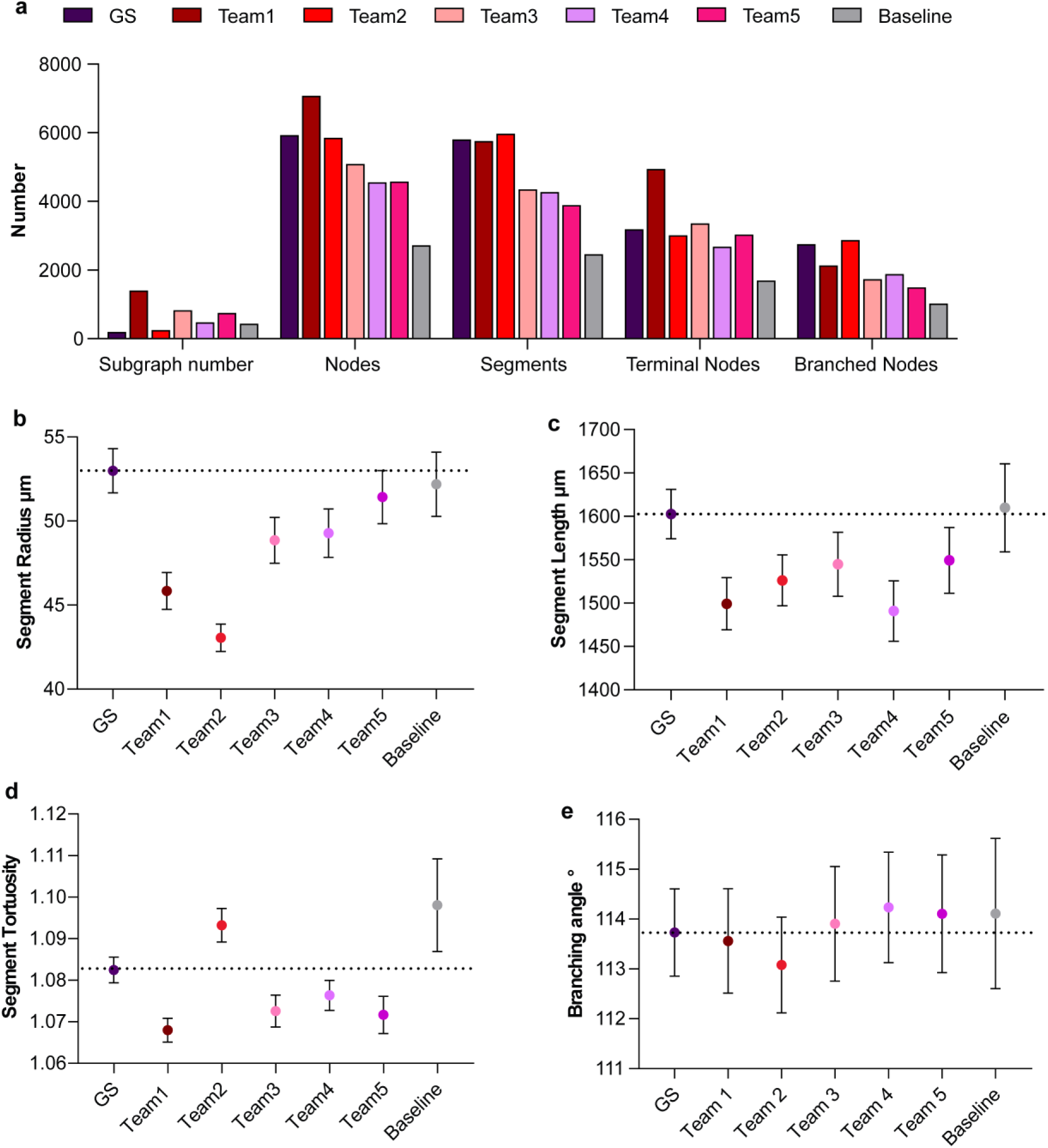
**a.** Bar chart showing the number of subgraphs, nodes, segments, terminal nodes, and branched nodes for all team solutions as well as the baseline model predictions and gold standard (GS) labels for private test data. **b-e.** Plots showing the radius, length, tortuosity of segments, and the branching angle between segments; mean and 95% Confidence Interval (CI) are shown for all metrics.

Team 1 has the highest number of segments, proportion of terminal nodes to branched nodes, and number of subgraphs, indicating the identification of many vessels but with a highly fragmented segmentation. Team 1 also has the second lowest mean radius indicating there is either an under segmentation of the vessel lumen or an increase in small vessels that may not be present in the gold standard.

Team 2 has the smallest number of subgraphs (247) comparable to that of the gold standard (199). It also has the most similar number of segments and proportion of terminal-to-branched nodes compared to the gold standard. This indicates better continuity in vessel segmentation, which can be attributed to the solution’s post-processing steps, and is also reflected in the team’s relatively higher clDice score. Team 2’s lower radius and length might also suggest under-segmentation or more conservative thresholding.

Team 3 serves as the midpoint of the teams having the third highest number of subgraphs behind Team 1 and Team 5, with a similar proportion of terminal-to-branched nodes as Team 5. By comparison to Team 5, Team 3 has a lower mean radius and length suggesting a more conservative thresholding strategy leading to a thinner radius and slightly better predictions for smaller unconnected vessels.

Team 4’s lower vessel length coupled with the relatively higher mean radius, lower number of subgraphs, and lower proportion of terminal-to-branched nodes, indicates a model which is more targeted to the larger well-connected structures in the network rather than the smaller peripheral vessels, and reflects the masking of small false positive structures in the background outside the kidney.

Team 5 has the largest average radius and length, the third highest number of subgraphs, and the second highest proportion of terminal-to-branched nodes. This implies that while the vessel thickness may have been more accurately predicted (from radius and length), many small unconnected structures were still identified leading to the high subgraphs and high proportion of terminal nodes. The team’s lower position on the LB suggests these small unconnected structures were not smaller unconnected vessel fragments but likely came from noise predicted outside the kidney, as seen in **Figure 3** and **Figure 4**.

## Discussion

Vasculature segmentation is a highly complex and challenging task in medical image segmentation due to the low proportion of vessel voxels, complex structure, and variable nature of human vasculature across different individuals. Citizen science enables experts and hobbyists alike from industry, academia, and government globally to engage in open and collaborative experimentation and development of solutions. Furthermore, all data and code are openly shared, serving as a benchmark for future algorithm performance evaluations and comparisons for HiP-CT data.

In this competition, participants employed various techniques to tackle the kidney vessel segmentation task. Notably, methods integrating data augmentation and self-training paradigms emerged as effective solutions to address challenges related to the small dataset. In addition to utilizing data augmentation, normalization and pseudo labeling techniques, implementing postprocessing steps to eliminate disconnected vessels proved essential for developing generalized solutions. Furthermore, some teams mitigated the challenges associated with highly unbalanced datasets by using custom loss functions, such as marching cube loss. The competition further highlights that the U-Net architecture^24^ remains a strong base model for medical image segmentation since all of the top five teams used variants of U-Net, customizing the ML models to enhance performance.

While this competition has generated interesting solutions for a fairly new imaging modality, several known limitations persist: (1) the models were trained on a relatively small dataset, which increases the risk of overfitting; (2) segmenting both the very fine and the large the microvascular network accurately still remains a challenge; (3) the competition might rely on specific evaluation metrics that do not fully capture the clinical relevance or quality of the segmentation, potentially overlooking important aspects of model performance in real-world applications; and (4) some models are computationally expensive and might be impractical or inefficient for deployment at scale.

Linking qualitative and quantitative interpretation of the teams’ outputs enables a more comprehensive understanding of ML model outputs and biases. It also allows assessment of the solutions in ways which are highly relevant for the potential downstream uses of such data. Based on the morphological analysis, a key challenge most teams faced was the high number of unconnected components and lower proportion of branched-to-terminal nodes compared to the gold standard. This has important implications particularly for modeling of blood flow or exosomes transport, as every terminal node requires a prescribed boundary condition, introducing a high dependence of model output onto the assigned boundary condition rather than the connectivity of the network as previously shown^18^. Simple post-processing strategies which remove unconnected components, such as that applied by Team 2, or more sophisticated approaches which seek to join fragmented vessel sections would be an important next step in utilizing the output of these models.

In future, the authors would like to further generalize the winning solution and train on other 3D HiP-CT datasets to explore the vasculature trees, exosome flow patterns, and morphological differences in organs other than the kidney.

## Methods

### Kaggle platform

The private leaderboard for identifying the five prize winners was finalized on February 8, 2024. The prize winners were awarded cash prizes (US$25,000 for first place; US$20,000 for second place; US$15,000 for third place; US$5,000 for fourth place; US$5,000 for fifth place). The teams submitted their inference code, after training their ML models, on the Submission portal. The submitted code was then run over the public test set to rank the teams on the public leaderboard. The teams typically use this score to validate their models. With each submission, the code is also run on the private test set for preliminary rankings on the private leaderboard (not visible to teams). Teams can make an unlimited number of submissions before the competition deadline but are limited to five submissions per day. On competition end, the teams can choose up to two solutions to submit as their final submissions, which are then scored on the private test set to rank the teams on the final private leaderboard. All scoring is done using the normalized surface dice as the evaluation metric and the top-5 teams on the final private leaderboard are selected as winners.

### Hierarchical Phase-Contrast Tomography data acquisition

HiP-CT is a propagation phase-contrast local tomography technique, as described by Walsh et al.^2^ Data acquisition follows the sample preparation and scanning protocols outlined in prior work^2,34^. Briefly, organs are fixed, partially dehydrated, and stabilized for tomographic scanning using an agar-ethanol mixture. HiP-CT scans are performed on one of two beamlines, BM05 or BM18 at the European Synchrotron Radiation Facility (ESRF). The scans are conducted hierarchically; the entire organ is first scanned at ca. 25 µm per voxel, followed by local tomography at specified locations within the intact organ at ca. 6.5 µm per voxel and ca. 1.3-2.5 µm per voxel.

Tomography scans are reconstructed into image volumes using a filtered back projection algorithm as detailed in prior work^34^. The final volumes consist of isotropic 3D image datasets, where higher resolution volumes of interest are registered within the larger organ volume using a rigid transformation. The contrast within these images arises from interference patterns caused by refraction of X-rays as they pass through the samples. Refraction is caused by physical density differences within the sample, and thus the modality particularly highlights edges between tissues with different densities.

### Gold Standard Label Acquisition

Five human kidneys were used to create the competition dataset. Three kidneys (Kidney 1, Kidney 2, and Kidney 3) were used for the training set. Two kidneys (Kidney 5 and Kidney 6) were used for the test set—Kidney 5 as the public test set and Kidney 6 as the private test set. Except Kidney 2, all were collected from donors who had consented to body donation to the Laboratoire d’Anatomie des Alpes Françaises prior to death. Kidney 2 was obtained after a clinical autopsy at the Hannover Institute of Pathology at Medizinische Hochschule, Hannover (Ethics vote no. 9621 BO K 2021). The transportation and imaging protocols received approval from the French Health Ministry. The basic scan parameters and demographic information of subjects can be found in **Supplementary Table 1**.

The segmentation of the five kidneys was conducted using Amira Version 2021.1. Initially, the raw image data underwent average binning, either x2 or x4. In x2 binning, the resolution was reduced from approximately 25µm to 50µm voxel size. In x4 binning, the resolution was reduced from 15.8µm to 63.1µm voxel size.

A 3D median filter was then applied using Amira-Avizo v2021.1 with 3 iterations and a 26 voxel neighborhood. To enhance the appearance of vessels, background detection correction was performed (Amira v2021.1; default settings). The segmentation process involved semi-manual techniques using Amira v2021.1’s magic wand tool, an interactive 3D region-growing tool. Curators selected a seed voxel within a vessel slice and refined parameters such as intensity threshold, contrast threshold, and hard-drawn limits to determine the stopping criteria for the 3D region growing. In cases where vessels were infilled with blood or largely collapsed, a manual voxel paintbrush tool was used to correct or fill the vessel in every slice. Further details and supplementary video of the semi-automated segmentation protocol is outlined in prior work^3^.

To ensure segmentation quality and provide error estimates on the gold standard, an expert segmentation validation process was implemented. An initial curator applied the above procedure on a dataset using three orthogonal views. An independent curator re-labeled the data adding to the labels of the initial curator, again using the three orthogonal planes. A third curator (referred to as proof-reader) was presented with 5-7 randomized 2D sections of the data in any one of the three orthogonal planes. They counted the number of vessel cross-sections in the slice. They recorded the number of true positive and false negative vessel cross-sections that were segmented. Note that a true positive means that the curators 1 and 2 have visually correctly labeled the vessel but does not assess the quality of the segmentation on a pixel-by-pixel basis. This ratio of true positive and false negative is used as the estimate for the number of vessels correctly segmented. The proof-reader then returns the data to the two curators, with the estimated segmentation proportion for each 2D area. The curators 1 and 2 then focus on 3D segmentation in areas where the proof-reading has highlighted the lowest vessel segmentation proportion. The process repeats iteratively until the proof-reader does not find any false negatives^3^. Note that false positives are not considered in the labeling/proofreading process as these are easily detected owing to their lack of connectivity to the main vessel tree. All data is made publicly available, see **Data Availability**.

### Evaluation metrics

The submitted solutions were ranked in the competition using Normalized Surface Dice (NSD) with tolerance (t) set to 0. NSD determines what fraction of segmentation boundary is predicted correctly. A boundary element is considered correctly predicted if the closest distance to the reference boundary is smaller than or equal to the specified threshold (tolerance). The tolerance determines the acceptable amount of deviation in pixels. The value of NSD lies between 0 and 1. The specific implementation^22,23^ proposed by Google Deepmind is used in the competition with an optimized version available as a notebook on the Kaggle platform at https://www.kaggle.com/code/metric/surface-dice-metric. To further evaluate the winning solutions and their performance post competition, four auxiliary metrics were chosen based on prior work by the authors^19^: Dice Similarity Coefficient (Dice)^35,36^, Centerline Dice (clDice)^37,38^, Average Symmetric Surface Distance (ASSD)^22,39^, and NSD (with tolerance=1). ASSD was computed using the MONAI library^40,41^ v1.3.0.

### Morphological analysis code

Vasculature in segmentation masks is reduced to spatial graph representations of the network through skeletonization. The resulting spatial graph describes the vessel network in terms of four entities: nodes, points, segments, and sub-segments. A segment is defined as being between a start node *i* and end node *j*; which correspond to either a branching point leading into another segment branch or a terminal end where no further branches were detectable. Nodes have an ID and a 3D spatial position (x, y, z). Between the start node *i* and terminal node *j* of each segment lie sub-segments with points, marking the start and end of each sub-segment able to capture the curvature of the segment. Each sub-segment has an associated radius *R_s_* and length *L*_s_. To create a spatial graph from a segmented image, we utilize an implementation of the parallelized version of the distanced ordered medial thinning algorithm^18,42^ in Amira v2021.2 termed the “Autoskeleton” plugin. The radius of each subsegment is estimated using 1/5th of the maximum Chamfer distance and the parameters applied are *smooth* = 0.5, *attach_to_data* = 0.25, and *iterations* = 10. We extract morphological parameters from the spatial graph following the definitions from prior work^18^. See **Code Availability**.

### Participation analysis

At the conclusion of the competition, participation metadata becomes publicly available on Meta Kaggle^43^—Kaggle’s public data on competitions, users, submission scores and kernels. This data is used to understand how the competition unfolded over its 3-month period. Analysis is implemented using standard python packages for data science such as Pandas, NumPy, Matplotlib, and Seaborn; creating all visualizations in Jupyter Notebooks, see **Code availability**.

### Statistical analysis

Kendall’s Tau^31–33^ (also called Kendall’s Rank Correlation) is used to quantify the agreement between two rankings and is independent of the number of entities ranked. *Tau* values closer to 1 mean a strong positive correlation between the two rankings—value of 1 means perfect alignment—whereas values closer to -1 mean a strong disagreement. A *p-value* associated with the *tau* value indicates the statistical significance of the correlation; lower *p-values* (closer to 0) indicate higher significance of the relationship between the two rankings such that it is unlikely to occur by chance. Kendall’s tau is computed using the implementation in the Python Scipy^44^ library.

## Acknowledgements

We extend our gratitude to the prize sponsors of this competition: Kaggle, Canadian Institute for Advanced Research (CIFAR), and Thermo Fisher Scientific. We thank Addison Howard, Ashley Chow, and Ryan Holbrook from the Kaggle Staff team for their expert support throughout the competition. We thank Heidi Schlehlein for assisting with the figures. This work is funded by the NIH Common Fund through the Office of Strategic Coordination/Office of the NIH Director under awards OT2OD033756 (K.B.) and OT2OD026671 (K.B.); by the Cellular Senescence Network (SenNet) Consortium through the Consortium Organization and Data Coordinating Center (CODCC) under award number U24CA268108 (K.B.); by the Kidney Precision Medicine Project grant U2CDK114886 (K.B.); by the NIDDK under awards U24DK135157 (K.B.) and U01DK133090 (K.B.); by grants CZIF2021-00642 (P.D.L & P.T), 2020-225394 (P.D.L & P.T) and 2022-316777 (P.D.L & P.T & C.L.W) from the Chan Zuckerberg Initiative DAF, an advised fund of Silicon Valley Community Foundation; ESRF funding proposals md1252 and md1290 (P.D.L), the Royal Academy of Engineering (CiET1819/10) (P.D.L), MRC (MR/R025673/1) (P.D.L) and T the NIH BRAIN Initiative CONNECTS program via National Institute Of Neurological Disorders And Stroke (NINDS) and National Institute Of Mental Health (NIMH) award UM1-NS132358 (C.L.W & P.D.L). The funders had no role in study design, data collection and analysis, decision to publish, or preparation of the manuscript. The content is solely the responsibility of the authors and does not necessarily represent the official views of the National Institutes of Health.

The authors sincerely thank those who donated their bodies to science so that anatomical research could be performed. Results from such research can potentially increase humanity’s overall knowledge that can then improve patient care. Therefore, these donors and their families deserve our highest gratitude.

## Data Availability

All training datasets are available on the competition website (https://www.kaggle.com/competitions/blood-vessel-segmentation/data). Additionally, complete competition dataset (including training datasets and test datasets), whole kidney datasets for test sets, team predictions, and trained model weights have been compiled into a single publicly available repository. The repository is made available as a Google Drive folder currently (https://drive.google.com/drive/folders/14hdA0JEuzdmBmLNik21WYLCmEtr6vSwv?usp=sharing), but the same will be submitted as a Zenodo Dataset before final publication. The raw imaging datasets are also available, and can be visualized, on the Human Organ Atlas data portal at https://human-organ-atlas.esrf.eu. The DOIs to individual datasets are available in the supplementary information table 1.

## Code Availability

All code for analysis and the winning solutions is publicly available on GitHub (https://github.com/cns-iu/hra-sennet-hoa-kaggle-2024). The code for morphological analysis is available on GitHub (https://github.com/HiPCTProject/Kaggle_skeleton_analyses).

## Author Contributions

Y.J. aided the design and implementation of the competition; contributed to the design and implementation of data analyses pre- and post-competition; contributed to the winning model validations; acted as a co-liaison to the Kaggle Staff team; co-wrote the paper. C.L.W. aided the design and implementation of the competition; contributed to the design and implementation of data analyses pre- and post-competition; acted as a co-liaison to the Kaggle Staff team; co-wrote the paper; assisted in collection of the HiP-CT datasets, contributed to gold standard segmentation labels, and co-developed the skeletonization pipeline and analysis. E.Y. assisted in segmentations; aided the design of the competition through baseline analyses and investigation of metrics; contributed to code validation and analysis of winning teams; co-wrote the paper. S.A. contributed to code validation on the winning teams; co-wrote the paper. S.N. contributed to code validation on the winning teams; performed the skeletonization analysis; co-wrote the paper. Y.Z. assisted in gold standard segmentations; contributed to code validation on the winning teams; co-wrote the paper. J.H. computed the auxiliary evaluation metrics; contributed to evaluation of solutions on the test dataset. K.S.G. aided the initial competition design. J.B. collected HiP-CT datasets; optimized imaging and reconstruction parameters. S.R. performed gold standard segmentations; assisted in data collection; co-developed skeletonization pipeline. P.T. collected HiP-CT datasets; optimized and performed HiP-CT image reconstruction and performed segmentations. A.B. sourced and prepared organs for imaging; provided histopathological expertise and training for gold standard segmentations. G.M.W. aided the competition design; manuscript revision. P.D.L. aided the conception, design, and implementation of the competition; revision of the manuscript. K.B. aided the conception, design and implementation of the competition; co-wrote the paper.

## Supplementary Notes

### Supplementary Note 1: Team 1 solution details

#### Summary

Team 1 proposed a customized U-Net^1^ based architecture model using a lightweight encoder named ConvNeXt-tiny^2^ with additional stem blocks^3^ to improve the feature extraction. A data augmentation technique, including random 3d slice rotation, was acquired to tackle the challenges related to the data. To make sure that the proposed model is optimized well, they trained the model with an ensemble loss combining focal loss^4^, dice loss^5^, boundary loss^6^ and a customized marching cube loss. The main goal of marching cube loss is to directly optimize the surface dice. The data were processed across three axes with large inference size and test time augmentation for inference.

#### Data Preparation

They trained the proposed model using all available data, including high-resolution and sparsely labeled data. Then, all slices were randomly resized to size 1536*1536. To increase the size of the training data, they sliced the whole volume along different axes and views. Heavy data augmentation, including flipping, scaling, adding noise, and intensity modifications, was utilized. Moreover, they randomly rotated the sampled slice in 3D space as online data augmentation.

#### Model

The general architecture of the model includes a 2.5D U-Net^1^ based architecture stacking 3 consecutive slices on channel dimension as input for the model to leverage 3D information. The encoder part of the model includes the ConvNeXt-tiny model^2^ combined with an extra stem block^3^ to extract high-resolution features. The decoder part of the model is a standard U-Net decoder. For the loss function, this team utilized a combination of four losses including focal loss^4^, dice loss^5^, boundary loss^6^, and a newly proposed customized Marching Cube loss.

Inspired by surface dice computation, the customized Marching Cube loss replaces the surface area expectation with the ground truth surface area. Therefore, the Marching Cube Loss (MCL) can be calculated as follows:

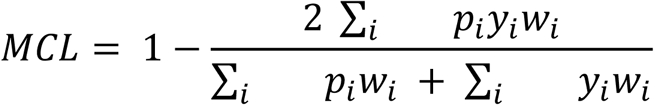

In practice, the surface area weight has minimal influence on the final performance of the mentioned loss. So, the loss mentioned above only considers the surface cubes (ground truth foreground and background cube’s surface areas are zero). Therefore, ignoring the surface area weight, this team used the mean of foreground, background and surface dice losses as the customized Marching Cube loss, which can be seen below.

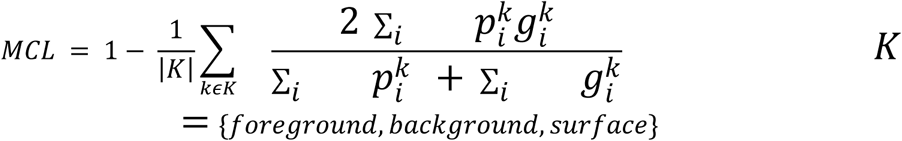

#### Training and Inference Details

This team trained the proposed model using a 32 batch size and a gradient accumulation process^7^. AdamW^8^ with additional WarmUpConsineAnnealing^9^ was used to optimize the model. The model was trained for 30 epochs.

In the testing process, all the slices were converted and resized to 3072*3072. The torch compile algorithm^7^ was utilized to accelerate the inference process. Moreover, the data were processed across three axes with large inference sizes and test time augmentation.

### Supplementary Note 2: Team 2 solution details

#### Summary

Team 2 used a customized 3D U-Net^10^ model for segmenting the kidney vessels. To overcome the generalizability problem, data augmentation techniques, including random positions and rotations, were used. Eventually, morphological post-processing techniques were used to remove small false positives and improve the model performance.

#### Model

The general architecture of the proposed neural network is based on 3D U-Net^10^. However, to overcome the problem of gradient vanishing, simple convolution layers were replaced by ResNet blocks^3^ in the encoder part of the model. To decrease the computational cost, all traditional up-convolutional layers have been replaced by up-sampling operations.

Online data augmentation techniques, specifically affine transformations such as rotation, scaling, and shearing, were employed to prevent overfitting. This process yielded a substantial increase in the number of patches, reaching over 7k for every epoch.

For the post-processing, the depth-first search algorithm^11^ was used to identify the unconnected, distantly placed chunks and remove them to increase the model’s overall performance.

#### Training and Inference Details

This team trained the proposed model with a batch size of 4 and a size of 128*128*32. For optimization, they used Focal loss^4^ and cosine decay scheduling^9^. The training lasted two weeks, and the maximum number of epochs was 700. 0.25 was used as the ultimate threshold for the inference to extract binary masks for vessels.

### Supplementary Note 3: Team 3 solution details

#### Summary

Team 3 employed a multifaceted approach to tackle the competition. They used a two-step process to refine sparse labels using dense labels from part of the training dataset. Initially, they trained two different UNet architectures (maxViT512^12^ and EfficientNetv2^13^) on densely labeled images to generate supplemental labels for sparsely labeled ones. These models were then further trained using the enhanced datasets, which included both dense and newly generated pseudo labels. The team trained three Unet models: EfficientNetv2, SeResNext101^14^, and MaxViT512, and one UNet++^15^ using all real labels from all kidneys plus pseudo labels.

#### Model

The main model, MaxVit512, represents a sophisticated UNet-based architecture combining CNN and Transformer methodologies, tailored specifically for medical image segmentation. It utilizes a hybrid multi-axis vision transformer mechanism, where both convolutional and self-attention mechanisms are employed at each stage of the decoder. This approach significantly enhances the model’s ability to distinguish between target objects and the background, crucial for effective segmentation. The MaxVit512 architecture, with its strategic use of a hybrid decoder, ensures high efficiency in segmentation with a reasonable computational and memory footprint. The models typically operate with a batch size of 32, allowing efficient processing of large datasets without compromising performance.

#### Training and Inference Details

Recognizing the variation in magnification between training and test sets (50µm/voxel for training and public test sets, and 63µm/voxel for the private test set), the team adjusted their training strategy to simulate the lower resolution of the private test set by scaling images down. They applied a ShiftScaleRotate augmentation, which randomly shifts, scales, and rotates the images to enhance model robustness to variations in image presentation. Training and inference were performed along different axes (x, y, and z) to manage varied resolutions and sizes, using solely 2D models. They maintained a consistent model resolution, opting for 512px for most tasks but switching to higher resolutions when supported by the dataset to minimize potential accuracy loss in smaller resolution data.

The team trained models on all available data once they observed stable convergence, maximizing learning from limited datasets. They used dynamic threshold values to maintain stability in model predictions, crucial for consistent segmentation performance. Given the intensity variations across different datasets, they implemented heavy intensity augmentation strategies to enhance model robustness. For final submissions, the team used both a single model and an ensemble of the four models mentioned, with both approaches yielding the same score of 0.727. Their script utilized CUDA for GPU acceleration, ensuring quick prediction processing. Through these strategic maneuvers, the team effectively managed dataset variabilities and optimized their models for high performance. Overall, MaxVit512 scored 0.727, and the ensemble submission also scored 0.727.

### Supplementary Note 4: Team 4 solution details

#### Summary

The fourth team’s final model strategy combined 2D and 3D models, utilizing d4 test-time augmentation (TTA)^16^ to improve segmentation results. They applied a multi-view TTA approach to the 2D models and conducted training using a 2-fold setup, selecting kidney_2 and kidney_3_dense as validation sets. They ensembled the models by assigning equal weights to both 2D and 3D models. Their training datasets included kidney_1_dense, kidney_2, kidney_3_dense, kidney_3_sparse, and pseudo labels. The team transitioned from slice-wise normalization to stack-wise normalization using percentiles to optimize model learning.

#### Model

The team’s main models included EfficientNet family models with a UnetPlusPlus decoder and SCSE attention^17^, integrated from the segmentation_models_pytorch library. EfficientNet-B5 was particularly effective for feature extraction, capturing complex patterns in images.

UnetPlusPlus enhanced feature propagation, crucial for accurately delineating boundaries and maintaining spatial details during encoding and decoding. Despite testing various architectures, the EfficientNet-B5-UNet++ model demonstrated superior performance. Their 3D model was based on the NN-Unet architecture but used the DynUnet from the MONAI library, with training over 500 epochs using SGD and a cosine annealing learning rate schedule.

#### Training and Inference Details

Training involved a multiview setup, stacking images in tensors and slicing along different axes. They employed a weighted sampling approach based on sample sparsity, with denser samples given a standard weight and sparser samples adjusted accordingly. For example, kidney_1_dense had a weight of 1, while kidney_2 was assigned a weight of 0.65. Dynamic augmentations like CutMix^18^ (applied with a 0.5 probability), shifts, flips, and brightness adjustments were crucial for model robustness. The 3D model’s augmentation strategy was simpler, involving d4 augmentations and random crops, with cropping adjusted to 192×192×192 to focus on volumetric capture.

Pseudo labeling was done using an ensemble of 2D models with data from https://human-organ-atlas.esrf.eu/, excluding some data to prevent leakage. Post-processing techniques, such as multiplying 3D model predictions with 2D model-derived ROI masks, significantly enhanced predictions. Ensembling predictions from 2D and 3D models with equal weighting further boosted performance. The team found that incorporating BoundaryDOULoss^19^ early in the competition significantly improved model performance, with a +2% increase in cross-validation metrics, +1.5% on the public leaderboard, and +5% on the private leaderboard.

### Supplementary Note 5: Team 5 solution details

#### Summary

Team 5 solution applied only a 2D segmentation model using HiP-CT slices as inputs. The final predictions were given by ensembling three 2D U-Net models trained on xy, yz and xz axes, respectively. The team trained the model on kidney_1, then kidney_2 and validated on kidney_3. To achieve better results, they applied several technical implementations such as pseudo-labeling from additional data, boundary loss and spatial resolution interpolation.

#### Pseudo-labelling

Considering that data is important to train an effective deep neural network, the team involved two additional datasets: LADAF-2020-31 kidney^20^ and LADAF-2020-27 spleen^21^, from the Human Organ Atlas (https://human-organ-atlas.esrf.eu/) into the training. A 4-step training scheme, was applied to gradually generate pseudo labels for the new datasets and incorporate them into training:

1. The team trained a base model on kidney_1 and pseudo-labeled, kidney_2;
2. Kidney_2 was incorporated into the training data together with kidney_1 to re-train the model. Then, they used this model to pseudo-labeled the LADAF-2020-31 kidney.
3. Similarly, they pseudo-labeled LADAF-2020-27 spleen dataset from the model re-trained on kidney_1, kidney_2 and LADAF-2020-31 kidney.
4. Finally, the model was trained on four datasets.

The 4-step training scheme enabled generating pseudo-labels for the new dataset. However, pseudo-labels can introduce training noise with false positives which are difficult to completely remove in this scheme without post-processing. To overcome this problem, the team used soft labels, the probabilities after sigmoid function, as pseudo-labels instead of hard labels.

#### Loss

As surface dice is one of the metrics in this competition, evaluating the predicted boundaries of the vasculature, the team applied a boundary weighted binary cross-entropy loss as equation:

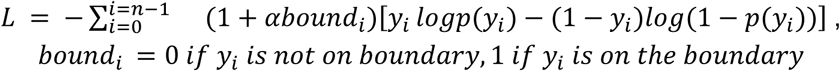

where n is the size of a mini-batch. α is a boundary weight and *bound_i_* is a binary value, indicating if the value *y_i_* is on the boundary. The last term is a typical binary cross-entropy function. The team set α as 0.9 to roughly double the loss on the boundary pixels. Apart from a boundary weighted binary cross-entropy loss, the overall loss function added dice loss^5^ and focal loss^4^.

#### Models

Four different 2D backbone networks: effnet_v2_s, effnet_v2_m, maxvit_base and dpn68 were ensembled for the final submission. They are implemented by a python package - Segmentation Models PyTorch (SMP)^22^. The input was cropped from xy, xz and yz axes in size of 512^2^, followed by pre-processing methods, such as 2D rotation, intensity contrast changes, horizontally and vertically flipped. After training, the inference was done with a sliding window method with an overlap size of half the input size.

#### Inference

Team 5 solution highlighted that 3D interpolation to make the voxel sizes of the training data and test data the same is important when performing vasculature segmentation inference on HiP-CT data. In this competition, the training data of kidney 1 to kidney 3 are ∼50 µm/voxel, however, the test dataset of kidney 6 is ∼63 µm/voxel. Team 5 applied a rescaling operation through 3 axes even though training was performed on 2D slices. The rescaling was implemented by trilinear interpolation and the rescaled data resulted in better performance.

### Supplementary Note 6: Useful strategies from other teams

#### Pre-processing

The kidneys were imaged by HiP-CT which requires the X-ray beam and scan setup to be independently tuned for every individual sample. Thus, scan parameters and hence the ranges of voxel intensities vary often dramatically between samples. This effect introduces weight bias and shifts when training deep neural networks. Therefore, several approaches were applied in this competition to construct a uniform input intensity space for stable training. Popular pre-processing techniques were normalization with global maximum and minimum across three training kidneys and intensity augmentation in training and validation. However, for 2D models, normalization of individual slices performs better than normalization with global maximum and minimum from 3D volume (https://www.kaggle.com/competitions/blood-vessel-segmentation/discussion/469022).

#### Model Selection

HiP-CT can image intact human organs, with isotropic voxels enabling clear visualization of 3D structures in any orientation. This inherently inspires the application of 3D as well as 2.5D and 2D segmentation models. Therefore, many teams explored different models for this task, reporting that 2D models were more efficient for achieving better results, due to the challenges of setting the hyper-parameters for 3D models. Due to limitations in computational resources for many competitors, 3D model input patches were smaller (input dimensions = 64 × 64 × 64) and thus contained less spatial context, compared to popular input settings of 512 × 512 in 2D and 2.5D models. If the input dimension were increased in 3D models, the batch size had to be reduced.

#### Post-processing

For HiP-CT scanning, organs are physically stabilized in cylindrical jars using an ethanol–agar or formalin-agar gel, so the images also contain the irrelevant background areas of the surrounding ethanol and agar. To remove the false positives on those areas, most of the teams created the mask for the kidney using different techniques such as Canny filters from OpenCV package and intensity thresholding. Since connectivity is one of the important factors for vascular trees and models failed in preserving these contextual long dependencies, connected component 3D (cc3d)^23^ was popularly used to improve the connectivity of the predictions as a post-processing method.

## Supplementary Figures

**Supplementary Figure 1.**
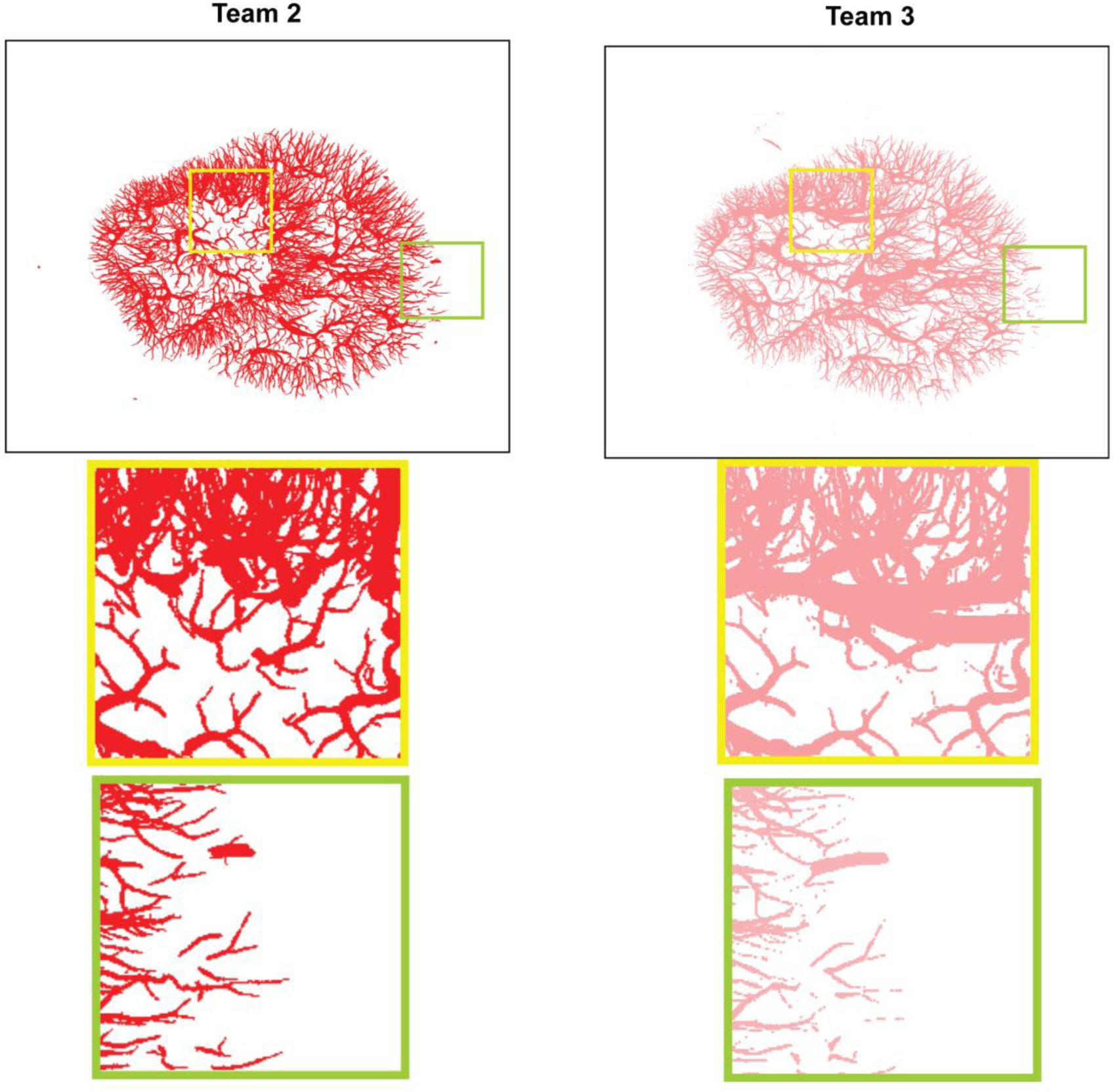
Maximum intensity projections (MIP) for Teams 2 and 3, with two insets per team in the yellow and green squares, respectively.

**Supplementary Figure 2.**
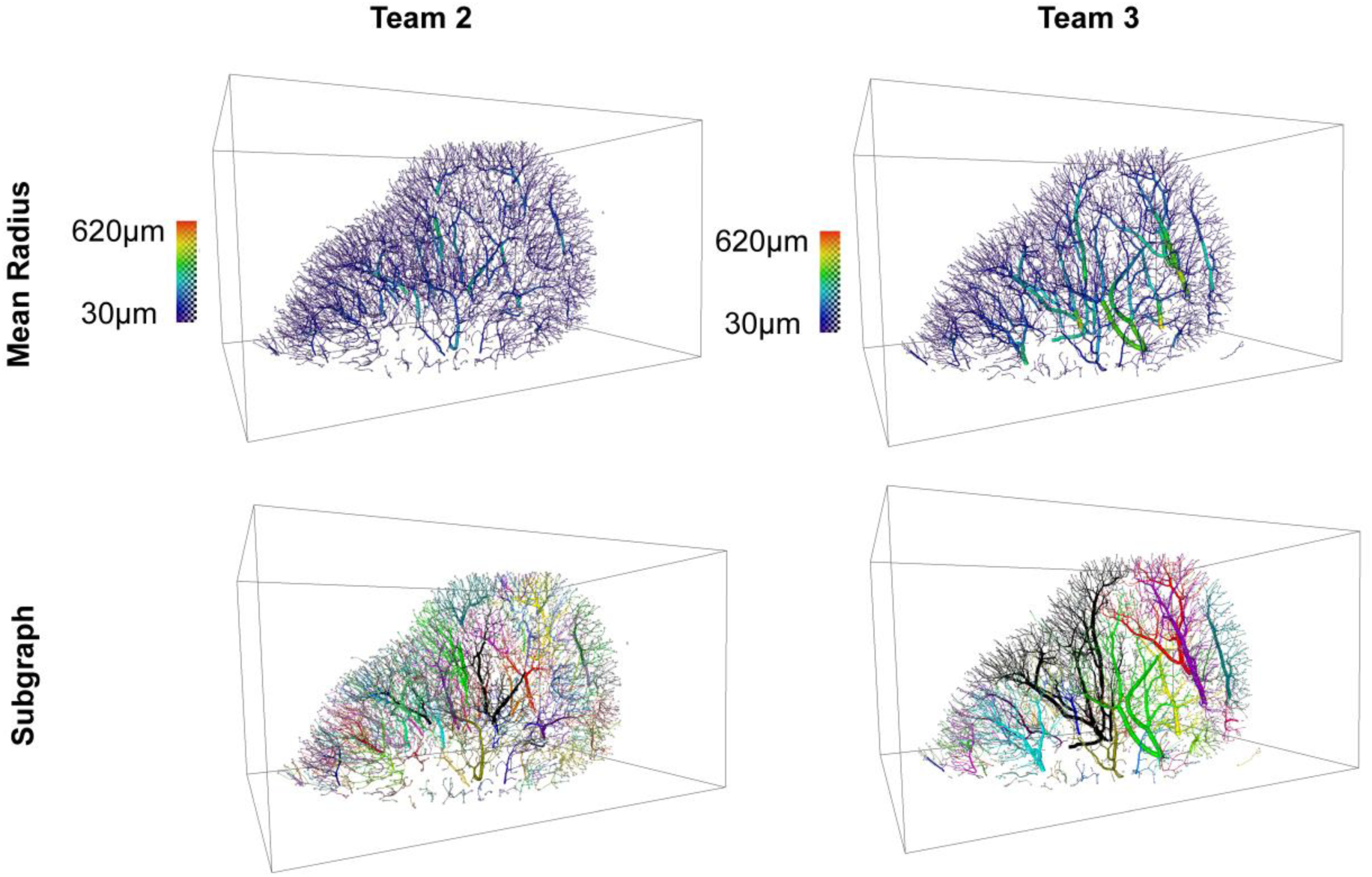
The figure shows the skeleton forms of the vessel network for Teams 2 and 3. In each case the vessel network size is proportional to the mean radius in the first row; the color shows the mean radius for each case. The second row is colored with each unconnected subgraph in a different color.

**Supplementary Figure 3.**
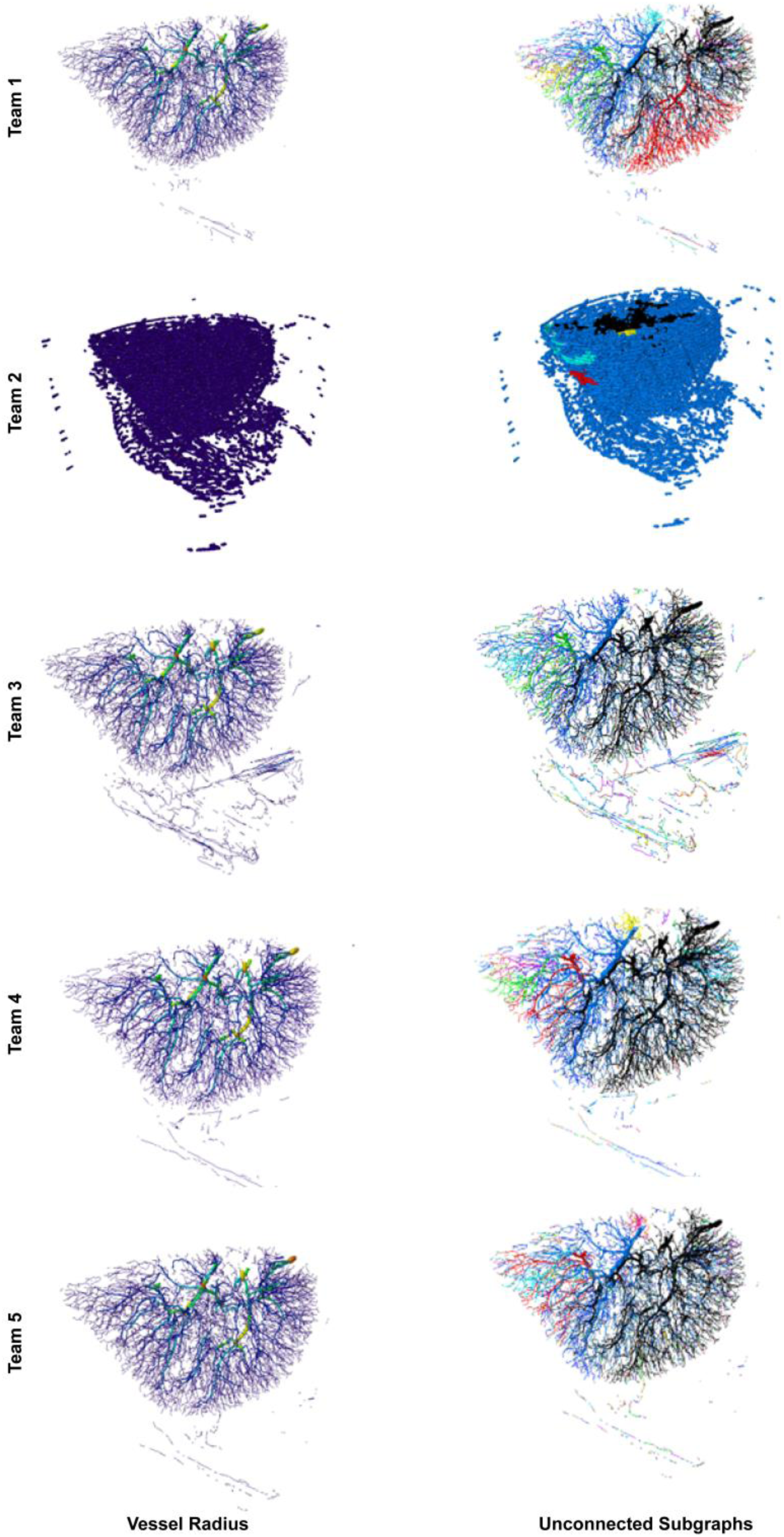
Showing the skeleton forms of the vessel network for all teams for the public test data. In each case the vessel network size is proportional to the mean radius in the first column; the color shows the mean radius for each case. The second column is colored with each unconnected subgraph in a different color.

**Supplementary Figure 4.**
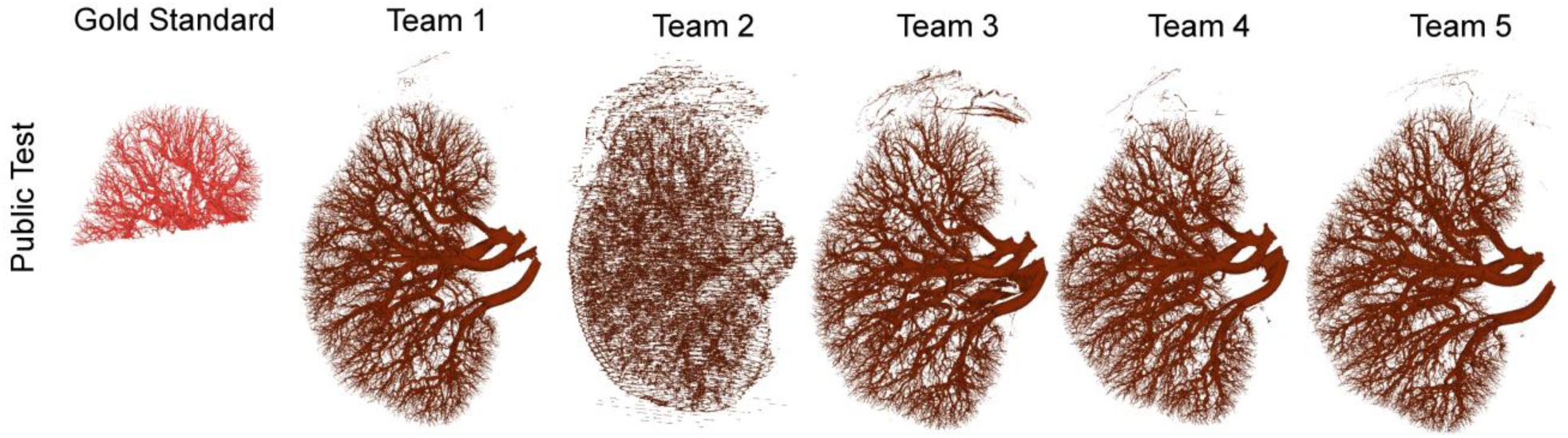
Visualization of the 3D output for inference of each team on the whole intact kidney datasets for the public test data (Kidney 5). Gold standard (red) shows the part of the whole kidney that was fully labeled and was part of the original competition dataset.

**Supplementary Figure 5.**
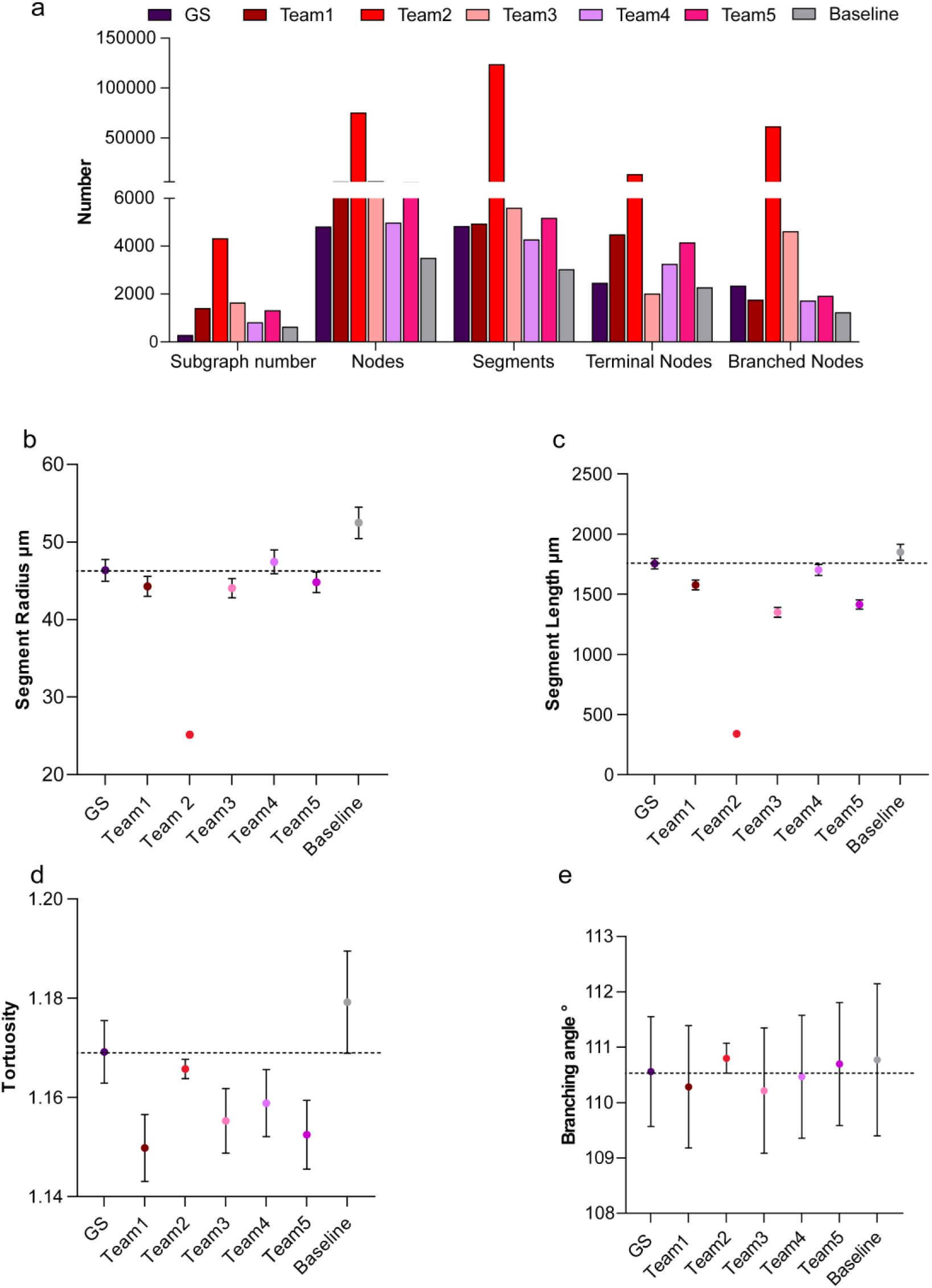
**a.** Bar chart showing the number of subgraphs, nodes, segments, terminal nodes, and branched nodes for all team solutions as well as the baseline model predictions and gold standard (GS) labels for public test data. **b-e.** Plots showing the radius, length, tortuosity of segments, and the branching angle between segments; mean and 95% Confidence Interval (CI) are shown for all metrics.

## Supplementary Tables

**Supplementary Table 1.** Listing of DOI and link for each dataset in the competition data.

| Kaggle identifier | Donor identifier | Dataset name | DOI | Visualization link |
| --- | --- | --- | --- | --- |
| 1 | LADAF-2021-17 | LADAF-2021-17_kidney_right_complete-organ_25.0um_bm05 | 10.15151/ESRF-DC-1773966439 | https://human-organ-atlas.esrf.eu/datasets/1773966586 |
| 1 | LADAF-2021-17 | LADAF-2021-17_kidney_right_VOI-03.1_2.6um_bm05 | TBC | link to visualization |
| 2 | S-20-28 | S-20-28_kidney_complete-organ_25.0um_bm05 | TBC | link to visualization |
| 3 | LADAF-2020-27 | LADAF-2020-27_kidney_left_complete-organ_25.08um_bm05 | 10.15151/ESRF-DC-572182553 | https://human-organ-atlas.esrf.eu/datasets/572182201 |
| 5 | LADAF-2021-17 | LADAF-2021-17_kidney_left_complete-organ_25.14um_bm05 | 10.15151/ESRF-DC-1773965419 | https://human-organ-atlas.esrf.eu/datasets/1773965003 |
| 6 | LADAF-2022-13 | LADAF-2022-13_kidney_2_complete-organ_15.77um_bm18 | TBC | TBC |

**Supplementary Table 2.**
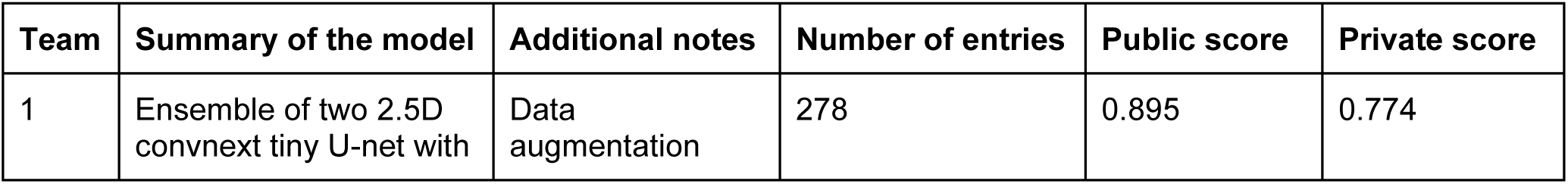
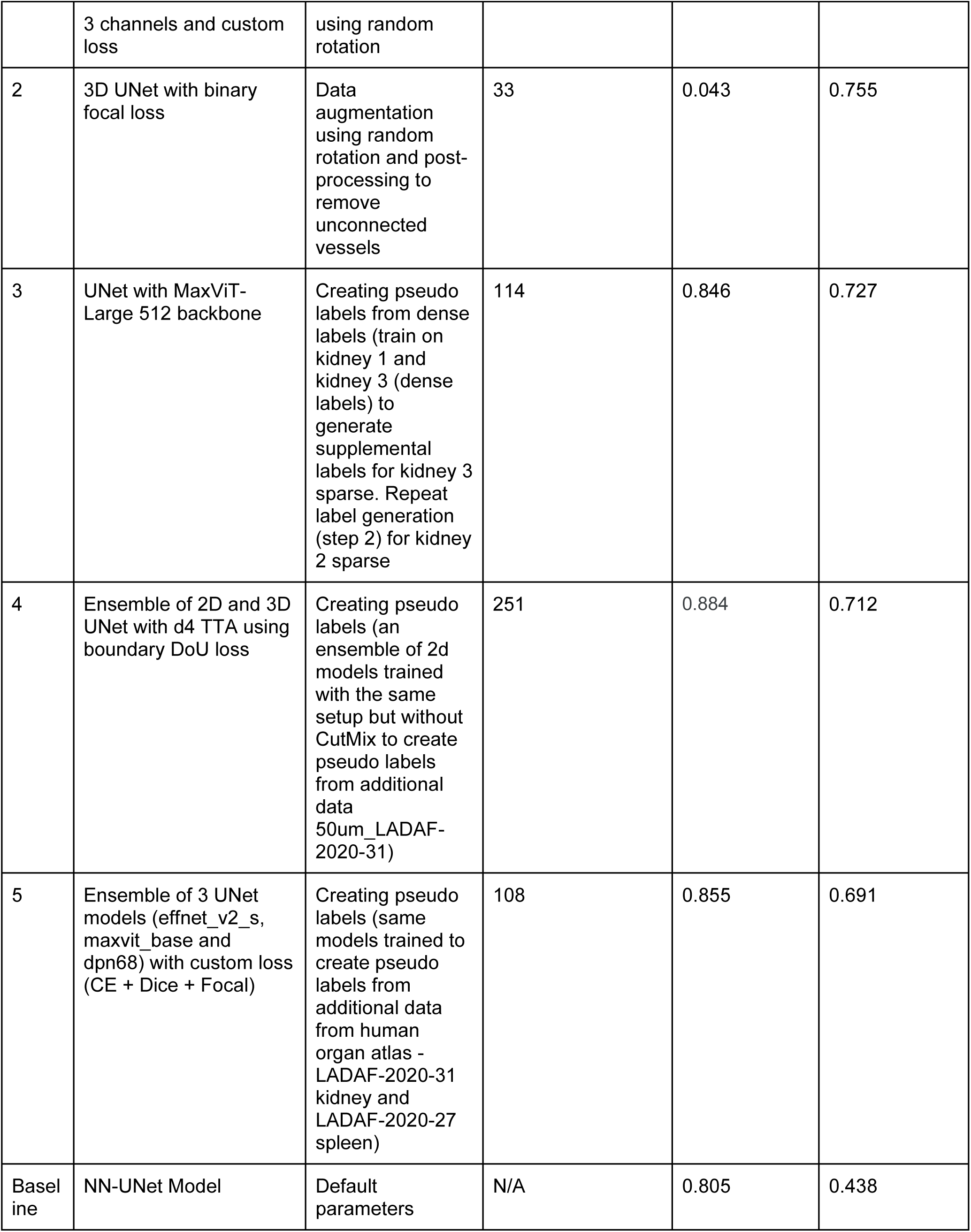
Scores for competition metric for top-5 teams and baseline model, including a brief summary.

**Supplementary Table 3.** All metric scores for top-50 teams, on both test sets. Team ID is based on ranking on the final private leaderboard. Lower is better for ASSD scores. Other metrics are bounded between 0 and 1, higher values are better.

| Team ID | Dataset ID | NSD (t=0) | NSD (t=1) | cIDice | ASSD | Dice |
| --- | --- | --- | --- | --- | --- | --- |
| 1 | kidney_5 | 0.8951 | 0.9569 | 0.833 | 1.3132 | 0.9283 |
| 1 | kidney_6 | 0.7741 | 0.9252 | 0.7859 | 0.7877 | 0.8289 |
| 2 | kidney_5 | 0.0431 | 0.0855 | 0.0134 | 40.5078 | 0.0119 |
| 2 | kidney_6 | 0.756 | 0.8479 | 0.7991 | 2.7445 | 0.6758 |
| 3 | kidney_5 | 0.8327 | 0.8959 | 0.7493 | 10.8552 | 0.8971 |
| 3 | kidney_6 | 0.7279 | 0.8844 | 0.7428 | 1.0919 | 0.8174 |
| 4 | kidney_5 | 0.8849 | 0.9423 | 0.8457 | 1.8925 | 0.9296 |
| 4 | kidney_6 | 0.7122 | 0.8623 | 0.7342 | 1.4183 | 0.8144 |
| 5 | kidney_5 | 0.8855 | 0.9554 | 0.8279 | 1.1549 | 0.9279 |
| 5 | kidney_6 | 0.6918 | 0.8612 | 0.7088 | 1.5988 | 0.8041 |
| 6 | kidney_5 | 0.8862 | 0.9477 | 0.8411 | 0.7425 | 0.9315 |
| 6 | kidney_6 | 0.6819 | 0.8055 | 0.6659 | 1.8331 | 0.792 |
| 7 | kidney_5 | 0.8594 | 0.9156 | 0.7959 | 6.1061 | 0.9152 |
| 7 | kidney_6 | 0.6768 | 0.8492 | 0.7139 | 1.4423 | 0.7802 |
| 8 | kidney_5 | 0.879 | 0.9431 | 0.8088 | 1.4752 | 0.9275 |
| 8 | kidney_6 | 0.6753 | 0.8585 | 0.7015 | 1.4384 | 0.778 |
| 9 | kidney_5 | 0.8272 | 0.8998 | 0.7227 | 4.2324 | 0.9088 |
| 9 | kidney_6 | 0.6651 | 0.8843 | 0.7343 | 1.1351 | 0.7758 |
| 10 | kidney_5 | 0.8103 | 0.8795 | 0.7159 | 10.2946 | 0.8912 |
| 10 | kidney_6 | 0.6572 | 0.8496 | 0.699 | 1.4754 | 0.7781 |
| 11 | kidney_5 | 0.6132 | 0.7349 | 0.6529 | 17.6852 | 0.649 |
| 11 | kidney_6 | 0.6559 | 0.7625 | 0.6824 | 3.2546 | 0.6951 |
| 12 | kidney_5 | 0.6362 | 0.8144 | 0.5687 | 7.2457 | 0.801 |
| 12 | kidney_6 | 0.6488 | 0.8012 | 0.6436 | 3.2204 | 0.7075 |
| 13 | kidney_5 | 0.8581 | 0.9165 | 0.7899 | 3.1011 | 0.9138 |
| 13 | kidney_6 | 0.6479 | 0.8382 | 0.6593 | 1.3991 | 0.7674 |
| 14 | kidney_5 | 0 | 0 |  | inf | 0 |
| 14 | kidney_6 | 0.6462 | 0.8177 | 0.6888 | 2.1497 | 0.7571 |
| 15 | kidney_5 | 0.8228 | 0.9023 | 0.7518 | 4.6402 | 0.9011 |
| 15 | kidney_6 | 0.6359 | 0.8184 | 0.6718 | 1.8454 | 0.7831 |
| 16 | kidney_5 | 0.8887 | 0.9503 | 0.8065 | 1.8009 | 0.9285 |
| 16 | kidney_6 | 0.6347 | 0.8635 | 0.6904 | 1.2871 | 0.7635 |
| 17 | kidney_5 | 0.8137 | 0.8884 | 0.6991 | 2.1368 | 0.9077 |
| 17 | kidney_6 | 0.6345 | 0.823 | 0.6739 | 1.8226 | 0.7782 |
| 18 | kidney_5 | 0.8368 | 0.8961 | 0.7846 | 7.3325 | 0.8789 |
| 18 | kidney_6 | 0.6245 | 0.8353 | 0.7032 | 5.0582 | 0.736 |
| 19 | kidney_5 | 0.8133 | 0.8869 | 0.6914 | 1.3084 | 0.9074 |
| 19 | kidney_6 | 0.6214 | 0.7885 | 0.6391 | 4.6257 | 0.724 |
| 20 | kidney_5 | 0.8568 | 0.9287 | 0.7776 | 2.5528 | 0.9155 |
| 20 | kidney_6 | 0.6167 | 0.8465 | 0.7333 | 1.4698 | 0.6847 |
| 21 | kidney_5 | 0.8199 | 0.891 | 0.7248 | 7.8449 | 0.896 |
| 21 | kidney_6 | 0.6141 | 0.8227 | 0.6838 | 2.1755 | 0.7271 |
| 22 | kidney_5 | 0.8722 | 0.943 | 0.802 | 1.5679 | 0.921 |
| 22 | kidney_6 | 0.6117 | 0.8155 | 0.6539 | 2.3056 | 0.7248 |
| 23 | kidney_5 | 0.8507 | 0.9086 | 0.7707 | 1.6365 | 0.9179 |
| 23 | kidney_6 | 0.6103 | 0.7729 | 0.654 | 3.0569 | 0.7025 |
| 24 | kidney_5 | 0.8735 | 0.9414 | 0.831 | 2.6179 | 0.9188 |
| 24 | kidney_6 | 0.6088 | 0.8202 | 0.6746 | 4.1682 | 0.7267 |
| 25 | kidney_5 | 0 | 0 |  | inf | 0 |
| 25 | kidney_6 | 0.608 | 0.8119 | 0.6585 | 1.9342 | 0.7667 |
| 26 | kidney_5 | 0.8676 | 0.9379 | 0.7936 | 1.3814 | 0.9224 |
| 26 | kidney_6 | 0.606 | 0.7712 | 0.6188 | 3.2132 | 0.7469 |
| 27 | kidney_5 | 0.8652 | 0.924 | 0.8019 | 5.6499 | 0.9108 |
| 27 | kidney_6 | 0.6035 | 0.7922 | 0.6504 | 2.6637 | 0.738 |
| 28 | kidney_5 | 0.8496 | 0.9096 | 0.77 | 5.6527 | 0.9124 |
| 28 | kidney_6 | 0.602 | 0.8053 | 0.6509 | 3.3284 | 0.7603 |
| 29 | kidney_5 | 0.8594 | 0.9187 | 0.7875 | 1.2754 | 0.9178 |
| 29 | kidney_6 | 0.5992 | 0.7264 | 0.594 | 3.1979 | 0.6882 |
| 30 | kidney_5 | 0.8703 | 0.9338 | 0.805 | 2.6242 | 0.9242 |
| 30 | kidney_6 | 0.5988 | 0.7907 | 0.6391 | 2.0989 | 0.7483 |
| 31 | kidney_5 | 0.8301 | 0.9013 | 0.7501 | 2.9544 | 0.9069 |
| 31 | kidney_6 | 0.5982 | 0.7992 | 0.6869 | 3.4409 | 0.7232 |
| 32 | kidney_5 | 0.8561 | 0.9326 | 0.7795 | 1.5592 | 0.9157 |
| 32 | kidney_6 | 0.5968 | 0.796 | 0.6446 | 2.1674 | 0.7349 |
| 33 | kidney_5 | 0.8432 | 0.9163 | 0.7432 | 3.3218 | 0.9129 |
| 33 | kidney_6 | 0.5959 | 0.7973 | 0.6252 | 6.5184 | 0.745 |
| 34 | kidney_5 | 0.8754 | 0.9397 | 0.8121 | 1.9118 | 0.9235 |
| 34 | kidney_6 | 0.5944 | 0.8322 | 0.6843 | 1.6091 | 0.7481 |
| 35 | kidney_5 | 0.8503 | 0.9269 | 0.764 | 2.2443 | 0.9123 |
| 35 | kidney_6 | 0.5943 | 0.7931 | 0.6531 | 3.0259 | 0.6511 |
| 36 | kidney_5 | 0.8952 | 0.956 | 0.8509 | 0.7754 | 0.9322 |
| 36 | kidney_6 | 0.5936 | 0.7696 | 0.609 | 2.7045 | 0.7683 |
| 37 | kidney_5 | 0.6082 | 0.7485 | 0.5749 | 24.0377 | 0.7656 |
| 37 | kidney_6 | 0.5905 | 0.7227 | 0.5769 | 6.9668 | 0.7079 |
| 38 | kidney_5 | 0.6804 | 0.7684 | 0.6423 | 19.6523 | 0.7369 |
| 38 | kidney_6 | 0.5905 | 0.7317 | 0.6036 | 5.5285 | 0.6497 |
| 39 | kidney_5 | 0 | 0 |  | inf | 0 |
| 39 | kidney_6 | 0.5896 | 0.8272 | 0.6714 | 1.733 | 0.7546 |
| 40 | kidney_5 | 0.7982 | 0.8653 | 0.7004 | 11.4582 | 0.8858 |
| 40 | kidney_6 | 0.5875 | 0.8007 | 0.639 | 2.9754 | 0.7554 |
| 41 | kidney_5 | 0.8348 | 0.9147 | 0.7845 | 4.2633 | 0.9016 |
| 41 | kidney_6 | 0.587 | 0.78 | 0.6293 | 4.6619 | 0.7572 |
| 42 | kidney_5 | 0.8811 | 0.9477 | 0.815 | 0.8389 | 0.9261 |
| 42 | kidney_6 | 0.5862 | 0.7822 | 0.6411 | 5.1143 | 0.6628 |
| 43 | kidney_5 | 0.8711 | 0.9382 | 0.7985 | 1.7559 | 0.9229 |
| 43 | kidney_6 | 0.5848 | 0.8085 | 0.6718 | 2.0455 | 0.6903 |
| 44 | kidney_5 | 0.8761 | 0.9428 | 0.8073 | 1.9085 | 0.9241 |
| 44 | kidney_6 | 0.5848 | 0.8102 | 0.6644 | 4.4716 | 0.6803 |
| 45 | kidney_5 | 0.8941 | 0.9541 | 0.8403 | 0.7352 | 0.9297 |
| 45 | kidney_6 | 0.584 | 0.827 | 0.6803 | 3.0314 | 0.6334 |
| 46 | kidney_5 | 0.8065 | 0.8823 | 0.6973 | 2.8374 | 0.9005 |
| 46 | kidney_6 | 0.5829 | 0.7626 | 0.6044 | 2.5261 | 0.7397 |
| 47 | kidney_5 | 0.8283 | 0.9073 | 0.7383 | 3.4346 | 0.905 |
| 47 | kidney_6 | 0.5824 | 0.7888 | 0.6594 | 2.2022 | 0.7001 |
| 48 | kidney_5 | 0.8741 | 0.942 | 0.8172 | 0.8616 | 0.9241 |
| 48 | kidney_6 | 0.5809 | 0.7819 | 0.6335 | 7.6572 | 0.7435 |
| 49 | kidney_5 | 0.7685 | 0.8545 | 0.715 | 12.7754 | 0.8565 |
| 49 | kidney_6 | 0.5788 | 0.738 | 0.6031 | 3.248 | 0.7292 |
| 50 | kidney_5 | 0.8364 | 0.9029 | 0.7445 | 3.5469 | 0.9101 |
| 50 | kidney_6 | 0.5782 | 0.799 | 0.6574 | 2.2052 | 0.7254 |

